# Soil heterogeneity and pleiotropy contribute to polygenic soil adaptation during postglacial range expansion in an alpine plant

**DOI:** 10.1101/2025.04.20.649708

**Authors:** Veronika Lipánová, Marco Griepentrog, Oliver Reutimann, Jonas Kussmann, Hirzi Luqman, Sebastian Dötterl, Simone Fior, Alex Widmer

## Abstract

1) Plants colonized new abiotic environments during postglacial range expansions. Little is known about whether populations adapt to different soil conditions during range expansion and, if so, which mechanisms underlie presumably polygenic adaptation. It remains unclear how pleiotropy and soil heterogeneity contribute to such adaptation.
2) We studied 43 populations of *Dianthus sylvestris* with characterized soil conditions along its postglacial range expansion in the Alps. We leveraged genome-wide data and variation in multiple soil variables to identify polygenic signatures of selection in soil-associated alleles using environmental association analysis, generalized dissimilarity models, and polygenic scores.
3) We found signatures of selection in 814 single nucleotide polymorphisms (SNPs) and the largest magnitude of allele frequency change in candidate SNPs associated with K, Mg, and Al. Candidate genes showed higher pleiotropy than randomly sampled genes. We found soil heterogeneity shaping the populations’ adaptive genetic variation in the landscape.
4) Our results suggest that populations of *D. sylvestris* adapted to contrasting soil chemical properties during postglacial range expansion through polygenic adaptation. Pleiotropy likely plays an important role in polygenic adaptation to novel selective pressures and soil heterogeneity is an important factor contributing to the maintenance of adaptive genetic variation.

## Introduction

Soil properties can be strong selective agents shaping plant adaptation, but they remain a largely underexplored dimension of the environment (Dauphin et al., 2022). Plants colonizing geologically heterogeneous landscapes, such as the Alps, face a variety of soil conditions. In a mosaic landscape formed by a diversity of geological materials, the weathering of rocks leads to the formation of soils with varying chemical and physical properties (Rajakaruna, 2018). Particularly in alpine regions, where soils are young and poorly developed (Poulenard & Podwojewski, 2006), variation in soil conditions can impose strong selection through metal toxicity and differences in nutrient and water availability. Strong selection can lead to the adaptation of populations to particular soil conditions (e.g. calcareous or siliceous), potentially limiting their ability to colonize contrasting soil conditions (Michalet *et al*., 2002; Parisod, 2022). Conversely, populations may adapt to the variety of soil conditions across soil boundaries, but this can come at the cost of reduced competitiveness in any specific environment (Rajakaruna, 2018).

Chemical soil properties are an important component of the abiotic environment. One of the key factors determining ion availability for plants is pH, which plays a crucial role in many soil processes and directly influences the solubility of soil compounds and elements (Binkley & Vitousek, 1989). For instance, strongly acidic soils (pH < 5) often contain an excess of toxic metals such as Al^3+^ and Mn^2+^ accompanied by a deficiency in essential nutrients like Ca^2+^ and Mg^2+^ (Foy, 1984; Kinraide *et al*., 2004). Specifically, soluble Al^3+^ represents the most limiting factor in acidic soils (Caniato *et al*., 2011; Long *et al*., 2024), as Aluminum is known to inhibit seed germination and root growth (Clarkson, 1965; Shen *et al*., 1993; Panda & Khan, 2009; Gould *et al*., 2014; Long *et al*., 2024). Manganese toxicity is also described as a limiting factor for plant growth in acidic soils (Foy, 1984; Fernando & Lynch, 2015). In contrast, neutral to alkaline soils (pH ≥ 7) typically have a relatively high levels of Ca^2+^ and Mg^2+^, while they are often deficient in the bioavailability of nutrients such as K and P.

Colonization of new environments during range expansions exposes populations to novel selective pressures, which can drive evolutionary changes in key traits (Barghi *et al*., 2020; Yeaman, 2022). In these new environments, genetic variants that confer an advantage are expected to increase in frequency, facilitating local adaptation (Barton & Turelli, 1989; Yeaman, 2015; Yeaman, 2022). Adaptive traits often have a polygenic architecture (Orr, 1998; Barghi *et al*., 2020; Bomblies & Peichel, 2022). Polygenic traits are expressed through combinations of many alleles of small effect, and genetic variation in a population provides ample material for selection to act on (Barrett & Schluter, 2008; Yeaman, 2015). Quantifying the cumulative effect of alleles associated with adaptive traits or environmental variables enables us to assess the adaptive genetic composition of populations shaped by selective pressures (Lehmann *et al*., 2023). Adaptation driven by polygenic traits can result in multilocus clines — patterns of allele frequency differences maintained by selection along environmental gradients and geographical locations (Barton, 1999; Barghi *et al*., 2020). We can study these changes in allele frequencies, i.e. allelic turnover, between populations by identifying genetic variation associated with e.g. soil elemental gradients (Fitzpatrick and Keller, 2015).

Understanding the mechanisms that maintain genetic variation in polygenic traits is fundamental to the study of evolution (Yeaman & Jarvis, 2006). During range expansions, a decrease in genetic variation in edge populations could hinder adaptation to novel environmental conditions. Conversely, increased genetic variation could enhance colonization success, promote long-term persistence, and facilitate adaptation to novel environments (Bridle & Vines, 2007; Barrett & Schluter, 2008; Crawford & Whitney, 2010). Heterogeneous landscape in environmental conditions provides substrate for spatially varying selection, a mechanism that contributes to the maintenance of genetic variation in environmentally associated loci (Byers, 2005; Yeaman & Jarvis, 2006; Delph & Kelly, 2014; Yeaman, 2015). Theory predicts that heterogeneous environments maintain higher levels of genetic variation, possibly leading to high variation in local adaptation, whereas, in homogeneous environments, most of the variation is purged (Levene, 1953; Huang *et al*., 2016; Booker, 2024). Spatially varying selection in heterogeneous environments leads to genetic differentiation among populations, i.e. to isolation by environment (IBE) – a positive correlation between genetic and environmental distances among populations, independent of geographic distance (Wang & Bradburd, 2014). Such heterogeneity is represented e.g. by population differences in soil chemical properties – soil heterogeneity (Vernham *et al*., 2023). Whether soil heterogeneity helps to maintain genetic variation and promotes adaptation to novel environmental conditions remains an open question.

Co-inheritance of traits, which can facilitate polygenic adaptation, is favored in heterogeneous environments with multiple selective pressures (Fisher, 1930; Orr, 2000). Pleiotropy, which occurs when a single gene affects multiple traits, is an important mechanism of trait co-inheritance. It has been shown that network connectivity can be used as a proxy for the degree of gene pleiotropy (Hamala *et al*., 2020; Rennison & Peichel, 2022). Empirical studies showed that pleiotropic genes are more connected and central within gene networks, thereby affecting the expression of other genes (Archambeault *et al*., 2020; Hamala *et al*., 2020; Rennison & Peichel, 2022; Williams *et al*., 2023; Whiting *et al*., 2024). Traditionally, pleiotropy has been considered as a limitation to adaptation, imposing ‘cost of complexity’ (Fisher, 1930; Orr, 2000). This is because if a mutation affects many traits, the likelihood of it having beneficial effects on all traits is assumed to be low. However, empirical studies suggest that the cost of complexity can be overcome by synergistic pleiotropy, with greater per-trait mutational effect sizes of pleiotropic genes facilitating rapid change of multiple traits at the same time (e.g. (Wagner *et al*., 2008; Wang *et al*., 2010; Frachon *et al*., 2017; Hamala *et al*., 2020)). It has been proposed that synergistic pleiotropy can be even a driving force of adaptive evolution (e.g. (Wagner & Zhang, 2011; McGee *et al*., 2016; Frachon *et al*., 2017)). Moreover, pleiotropic genetic changes can be favoured by selection when populations colonize new environmental conditions, because they enable larger steps to new trait optimum (Wagner *et al*., 2008). Range expansion, when species colonize new environments, can represent such a scenario of sudden shifts in optimal trait values (Des Marais *et al*., 2017; Hamala *et al*., 2020). However, the extent to which pleiotropy is involved in adaptation remains largely unknown.

One of the species that experienced range expansion in response to Quaternary climatic fluctuations is *Dianthus sylvestris*. While postglacial dynamics of adaptation to different climate conditions have been studied (Luqman *et al*., 2023), adaptation to the highly heterogeneous soil conditions that form a mosaic across the Alps, has yet to be explored. Luqman et al. (2023) identified glacial refugium around Monte Baldo and the western Dolomites, characterized by calcareous-like conditions. From that area, the species expanded across the Alps, colonizing environments heterogeneous in chemical soil properties. Further, postglacial recolonization of the Alps from glacial refugium led to loss of the genetic diversity in populations along range expansion (Luqman *et al*., 2023). Whether this loss of genetic variation also affected adaptive genetic variation (i.e. variation associated with soil variables) and if soil heterogeneity contributed to the maintenance of adaptive genetic variation remains unclear.

Here, we study the polygenic basis of adaptation to contrasting soil environments along the route of postglacial range expansion and how mechanisms such as pleiotropy and soil heterogeneity contribute to or constrain such polygenic adaptation. Using genome-wide sequence data of 43 populations of *Dianthus sylvestris* sampled across the Alpine arc, we asked: (i) Are there genetic signatures of selection in *D. sylvestris* populations from contrasting soil environments? (ii) How do soil variables drive selection and shape the adaptive genetic composition of populations? (iii) Which genes are under selection and contribute to soil adaptation? (vi) Are candidate genes of higher pleiotropy than randomly sampled sets of genes? (v) Did the loss of genetic variation during range expansion affect adaptive genetic variation and did the soil heterogeneity contribute to the maintenance of adaptive genetic variation in populations? To answer these questions, we applied approaches linking environmental and genetic data and revealed how the frequencies of candidate alleles vary in heterogeneous soil environments along the range expansion. We assessed the adaptive genetic composition of populations in response to the main drivers of soil adaptation in *Dianthus sylvestris*. Further, we inspected the positions of candidate genes in the gene networks, their interactions with other genes, and tissue specificity in gene expression as a proxy for pleiotropy. Lastly, we explored the relationship between populations’ adaptive genetic variation and soil heterogeneity.

## Materials and Methods

### Field soil sampling

In 2023, we sampled soil in each population locality (in total 43 populations) from five sub-sites to cover site spatial variation in chemical soil properties. We first removed the organic layer (the upper layer up to ∼5 cm below ground), then we took the soil sample from the rhizosphere, proximity of roots, ∼5-15 cm below ground. Soil samples were air-dried afterward, sieved through a 2 mm sieve, and five sub-sites samples/population were mixed to get one composite soil sample/population.

### Soil pH and exchangeable cations measurements

To characterize soil chemical properties, we measured soil pH and exchangeable cations. Soil pH was determined in 0.01 M CaCl2 solution (soil:solution ratio of 1:5). After shaking for 10 min, the samples were left to rest for 24 hr before measurement in suspension using a pH meter (713 pH Meter, Metrohm, Switzerland). Exchangeable cations (Al, Ca, Co, Fe, K, Mg, Mn, Na, P, and Sr) were extracted with 0.1 M BaCl2 (soil:solution ratio of 1:6.5) for 2 hr on a horizontal shaker, filtered through paper filters (Whatman 42) and diluted 1:10 prior to quantification with inductively coupled plasma – optical emission spectroscopy (5100 ICP-OES, Agilent Technologies, USA) (Hendershot & Duquette, 1986). The analytical lines used were Al 396.152nm, Ca 422.672nm, Co 228.615nm, Fe 234.350 nm, K 766.491nm, Mg 285.213nm, Mn 260.568nm, Na 588.995nm, P 213.618nm and Sr 216.596nm. The operation conditions of the ICP-OES analyses were as follows: sample flow rate 1.5 mL min−1, RF power 1.2 kW; plasma, auxiliary and nebulizer gas flow rates 12.0, 1.00, and 0.70 L min−1, respectively, view height 8.0mm, three replicated reading on-peak 5 s. A multielement standard (1000 mg/l) containing Al, Ca, Co, Fe, K, Mg, Mn, Na and Sr as well as a single P standard (1000 mg/l) were used for instrument calibration. External calibration standards were prepared for 0.1, 0.2, 1, 2, 10, 20, 100 mg/l. Additionally, Scandium (Sc) was used as internal standard to correct for matrix effects. Therefore, all calibration standards and all samples were spiked with Sc to obtain a concentration of 2.5 mg/l in all solutions and Sc was measured on analytical line 361.383nm. Sample concentrations were calculated using external calibration method within the instrument software.

### Population genomic data and inference of population genetic structure

We leveraged available whole-genome sequencing data (∼ 2x mean coverage) of *Dianthus sylvestris* populations for 43 populations (14 individuals/population) covering the species’ distribution and representing the post-glacial colonization of different soil environments across the Alps (Luqman *et al*., 2023). Briefly, variants were identified using freebayes v.1.3.1 (Garrison & Marth, 2012) and population allele frequencies were calculated based on genotype likelihoods via vcflib popStats v.1.0.1.1 (Garrison *et al*., 2022). For further details see Luqman et al. (2023).

We inferred the genetic structure of 43 populations using principal component analysis (PCA) based on whole genome population allele frequencies (∼ 10 million SNPs). The analysis was conducted with the ‘fviz_pca_ind’ function from the ‘factoextra v.1.0.7’ R package (Kassambara & Mund, 2020).

### Statistical analyses of soil data and genetic variation partitioning

To explore the variation among populations in the soil variables (pH and exchangeable Al, Ca, Co, Fe, K, Mg, Mn, Na, P, and Sr; Data S1), we analysed PCA biplot using the ‘fviz_pca_biplot’ function via the ‘factoextra v.1.0.7’ R package (Kassambara & Mund, 2020). Further, we calculated pairwise Spearman’s correlation coefficients using the ‘corrplot’ function from the R package ‘corrplot v.0.92’ (Wei & Simko, 2024) to inspect the relationship among soil variables. To further explore the similarity of populations in soil variables (population clustering into ‘soil environments’), we used the ‘pheatmap’ function from the R package ‘pheatmap v.1.0.12’ (Kolde, 2019) to aggregate populations using complete-linkage method of hierarchical clustering. We applied min-max normalization of the variables and scaled them to range between −1 to 1.

We applied partial redundancy analysis (pRDA, (Capblancq & Forester, 2021)) to disentangle drivers of selection across the Alps. Specifically, we tested what proportion of variance in the genetic data can be explained by soil variables, after accounting for population genetic structure. We ran pRDA models using the ‘rda’ function from the R package ‘vegan v.2.6.4’ (Oksanen, 2022) with nine soil variables as explanatory variables, excluding Fe and Mn due to at least one absolute pairwise Spearman’s correlation coefficient exceeding 0.7 (Dormann *et al*., 2013). We accounted for the effect of population structure by including principal component 1 (PC1) scores extracted from PCA based on genetic data in the pRDA. We used population allele frequencies as response variables in all the models. We tested the following models: (i) full model – both soil variables and population genetic structure as explanatory variables, (ii) soil variables model – soil variables as explanatory variables and population genetic structure as a conditioning variable, and (iii) population genetic structure model – population genetic structure as explanatory variables and soil variables as conditioning variables (overview of models in Table S1a). Lastly, we included climatic variable models in addition to the listed models above. We obtained monthly climatic variables with a spatial resolution of approximately 650 x 650 m for the study area (Dauphin *et al*., 2021) from the CHELSA v.2.1 database (Karger *et al*., 2017). We ran another set of pRDA models including also eight uncorrelated climatic variables (see Data S1 for a complete overview of variables) for comparison of explained variance in genetic data by different sets of environmental variables (overview of models in Table S1b).

### Environmental association analyses

We performed environmental association analyses using latent factor mixed models (LFMM2, (Caye *et al*., 2019)) to identify population allele frequencies that are significantly associated with soil variables. We ran a univariate environmental association analysis with each of the 11 soil variables (Al, Ca, Co, Fe, K, Mg, Mn, Na, P, Sr, and pH) as explanatory variable and population allele frequencies (whole genome SNP dataset; ∼10 million SNPs) as response variable. We accounted for neutral genetic structure by latent factors (*K*). We applied the ‘lfmm_ridge’ function from the R package ‘lfmm v.1.1’ (Jumentier *et al*., 2022) to estimate the effect of the environmental variable on allele frequencies using the regularized least-squares problem ‘ridge estimates’. We chose *K* = 4 as the number of latent factors accounting for population structure. For the choice of the *K*, we followed Cattell’s rule, retaining principal components corresponding to eigenvalues up to the point of inflection in the scree plot where the explained variance decreases (Cattell, 1966; Luu *et al*., 2017) (Figure S1). For significance testing, we corrected for multiple testing by transforming *p*-values (calibrated already by genomic inflation factor) to *q*-values using the ‘qvalue’ function from the R package ‘qvalue v.2.3’ (Storey *et al*., 2022) and applied a false discovery rate (FDR) of 0.05. To refine the list of significantly associated SNPs (Data S2), we applied the following steps: (i) retaining only SNPs in genic regions (annotated draft genome assembly available at https://doi.org/10.5061/dryad.x0k6djhng), (ii) filtering out low-frequency (< 5%) and high-frequency alleles (>95%), (iii) retaining only genes with at least two significantly associated SNPs (to reduce possible false positives), and finally, (iv) selecting one SNP with the highest effect size (estimated in LFMM2 and representing magnitude of the allele frequency change) for each gene (in case of multiple SNPs associated with different soil variables in one gene, we kept all of these). The final set of SNPs from LFMM2 is called soil-associated SNPs (Data S3). For an overview of the number of SNPs in each filtration step see Table S2.

### Generalized dissimilarity modelling

We examined which soil variables were associated with the largest magnitude of allele frequency change (allelic turnover) in SNPs and how rate of change in allele frequencies varied along particular soil elemental gradients (variation in exchangeable cations of particular element among populations) (Figure S2). To do so, we applied a non-linear distance-based approach – generalized dissimilarity models (GDM, (Fitzpatrick & Keller, 2015)) using the R package ‘gdm v.1.5’ (Fitzpatrick *et al*., 2022). This method uses I-spline functions for fitting the relationship between population dissimilarity and environmental variables as predictors (Ferrier *et al*., 2007). We adopted an approach investigating separately each soil-associated SNP, identified by LFMM2 (Fitzpatrick & Keller, 2015; Dudaniec *et al*., 2018; Tóth *et al*., 2023; Van Deurs *et al*., 2025). We calculated the genetic distance between all possible pairwise population combinations for each soil-associated SNP as Nei’s pairwise *F*_ST_ (Nei, 1987) using the ‘pairwise.neifst’ function from the R package ‘hierfstat v.0.5’ (Goudet & Jombart, 2022) and scaled values to range between 0 and 1 by subtracting the minimum value and then dividing by the maximum minus the minimum value, as suggested in (Fitzpatrick & Keller, 2015; Dudaniec *et al*., 2018). Changes in genetic distance between populations along soil elemental gradient represent allelic turnover. We extracted partial allelic turnover for each variable per each SNP corresponding to the relative importance of the predictor to the total allelic turnover (Fitzpatrick & Keller, 2015). In addition to soil variables (Al, Ca, Co, Fe, K, Mg, Mn, Na, P, Sr, and pH) as predictors in the multivariate models, we included also Euclidean geographic distance to test whether allelic turnover along soil elemental gradient was better explained by geographic distance than by changes in exchangeable cations. We removed 78 SNPs, for which there was no significant allelic turnover associated with any of the soil variables or geographic distance in tested models as suggested by (Van Deurs *et al*., 2025). We estimated the relative importance of each variable by ranking the magnitude of allelic turnover (i.e. changes in *F*_ST_ along the gradient of specific variables) relative to the magnitude of allelic turnover of another variable within each tested SNP. For each SNP, we also extracted deviance (goodness of the model fit) explained by the GDM. Further, we removed SNPs with a higher magnitude of allelic turnover in response to geographic distance than to soil variables (Figure S3a, b). Afterwards, as we still observed relatively high importance of geography (Figure S3c), we kept only SNPs with the lowest association with geographic distance (relative importance <= 0.047; 25% of the distribution of relative importance values; removing in total 2933 SNPs). According to the relative importance of variables, we assigned candidate SNPs as associated with specific elements (compared to LFMM2, 21% of associations were reclassified, see Data S4). Finally, we retained for each soil variable per each gene (consistent with the filtering procedure of soil-associated SNPs from LFMM) one SNP with the highest variance explained by GDM resulting in a final list of candidate SNPs (Data S4). To interpret the turnover functions, we inferred the maximum height of the GDM fitted spline indicating the allelic turnover, i.e. magnitude of allele frequency change. Magnitude of allele frequency change also assesses the importance of the ecological variable in explaining allelic turnover at that locus (Figure S2; (Fitzpatrick & Keller, 2015). Further, the shape of the GDM fitted spline reveals how the rate of change in allele frequencies varies along the soil elemental gradient (Fitzpatrick & Keller, 2015). For instance, steep slope of the spline indicates strong response and flat slope of the spline indicates weak response at the particular part of the soil elemental gradient. Furthermore, we calculated the mean and standard deviation of partial allelic turnover for each category of candidate SNPs associated with a specific element and we plotted the allelic turnover functions of each candidate SNP.

### Polygenic scores

We assessed the adaptive genetic composition of populations in many candidate loci based on polygenic scores (PS), while accounting for their varying effect sizes. Applying this approach, we can identify locally adaptive traits in populations using genomic data and assess how the polygenic scores correlate with environmental variation (Beer *et al*., 2022; Wadgymar *et al*., 2022; Folkertsma *et al*., 2024; Van Deurs *et al*., 2025). We calculated PS for each population based on candidate SNPs associated with the exchangeable cations that showed the highest magnitude of allelic turnover (i.e. K, Mg, and Al). Separately for each set of candidate SNPs, we summed up allele frequencies at particular SNP multiplied by standardized effect size (extracted from LFMM2 with standardized exchangeable cations). Using standardized effect sizes allowed us to compare effects of different soil variables. We explored the relationship between PS with either exchangeable K, Mg, or Al using the ‘gam’ function from the R package ‘mgcv v.1.9’ (Wood, 2011).

Specifically, we investigated the associations with exchangeable Al, as high Al levels are toxic for plants and can impact their fitness. We asked whether genetic composition of populations based on Al-associated SNPs, can predict the germination rates of different populations in the selective treatment – soils with high Al levels. We conducted a germination experiment with six populations (128 seeds/population, i.e. 768 seeds in total, see Data S5 for an overview of experimental populations) and two soil treatments of high exchangeable Al (17 mg/kg and 24 mg/kg). To stimulate germination, we placed Petri dishes with the seeds sown in soil prior the experiment into an incubator under 4°C for 10 days. After this period, we exposed Petri dishes to the conditions of 12 hours daylight at 20°C and 12 hours dark at 18°C and we recorded the germination as the appearance of cotyledon leaves for the period of 85 days. We tested the effect of population cumulative genetic composition (by polygenic scores) on germination using the ‘glmer’ function, binomial generalized linear mixed effect regression model, from the R package ‘lme4 v.1.1’ (Bates *et al*., 2015), accounting for soil treatment and Petri dish as a random factor. Further, we predicted for each population the probability of seed germination based on the output from glmer models and polygenic scores using the function ‘predict_response’ (type = ‘random’, interval = ‘confidence’), predictions were corrected for population level and soil treatment effects; R package ‘ggeffects v.1.7’ (Lüdecke, 2018).

### Identification of *Arabidopsis thaliana* orthologs, Gene ontology enrichment

For further functional interpretations of candidate genes, gene ontology (GO) enrichment, and proxies of pleiotropy (gene-gene interactions, tissue specificity, and centrality statistics), we identified *Arabidopsis thaliana* orthologs by reciprocal best blast hit (RBBH) between protein sequences of *D. sylvestris* and *A. thaliana* (TAIR10). We used blastp (Camacho *et al*., 2009) in the RBBH analysis to identify the best matching protein sequences between the two species by using each set of protein sequences as both the query and the subject (‘database’). To reduce multiple hits, we applied the following filtration criteria of keeping only the hits with the highest bitscore and the lowest e-value. Lastly, we applied a strict threshold of at least 70% identity in the alignment for the final refinement of the candidate gene list (Data S6). We extracted the description of *A. thaliana* orthologs from Araport11 annotation (Araport11_GFF3_genes_transposons.current.gff.gz; last update 10/2024) available in TAIR database (www.arabidopsis.org).

For evaluation of functional importance of the candidate genes, we performed a gene ontology (GO) enrichment analysis within biological processes (BP) and molecular functions (MF). We applied Fisher’s exact test (p < 0.05) with the ‘elim’ algorithm implemented in Bioconductor package ‘topGO v.2.5’ (Alexa & Rahnenführer, 2022). *Arabidopsis thaliana* was used as the background gene set in all analyses. We built the database using Bioconductor package ‘biomaRt v.2.54’ (Durinck *et al*., 2005). The ‘elim’ algorithm traverses the GO hierarchy from the bottom to the top, discarding genes that have already been mapped to significant child terms while accounting for the total number of genes annotated in the GO term (Grossmann *et al*., 2007; Alexa & Rahnenführer, 2009). We excluded GO terms with more than 500 genes as broad GO categories do not provide information about specific processes and functions (Bohutínská *et al*., 2021) (Data S7 overview of all GO terms). We also excluded GO terms with only one associated gene. We calculated the fold enrichment (FE) for each significantly enriched GO term by dividing the number of significant genes in our dataset annotated to that particular GO term by the expected number of *A. thaliana* genes annotated to that term based on the background distribution (Data S7). Lastly, we plotted a selection of significantly enriched GO terms relevant to soil adaptation (Data S7; manual search for GO terms related to drought – stomata development; root development; ion transport, and ion homeostasis).

### Quantification of pleiotropy

We applied a similar approach to (Nocchi *et al*., 2024; Whiting *et al*., 2024) to quantify pleiotropy. We applied phenotype-free approach and used proxies for pleiotropy such as gene network connectivity – centrality statistics and number of gene-gene interactions (ATTED-II database, (Obayashi *et al*., 2022); STRING database (Szklarczyk *et al*., 2015)). Further, we explored tissue specificity of gene expression leveraging gene expression data for *A. thaliana* from Expression Atlas (Papatheodorou *et al*., 2018). To assess gene network connectivity, we applied the following centrality measures: node closeness, node degree, node strength, and node betweenness. Specifically, node closeness measures a node’s ability to interact with all other nodes, including those not directly connected, thereby indicating its co-expression potential across the gene network. Node degree represents the number of nodes connected to a gene, indicating the number of co-expressed genes. Node strength reflects the total weight of these connections. Node betweenness captures a node’s role as a bridge within the network, connecting different co-expression sub-networks. Each of these metrics is expected to positively correlate with pleiotropy (Hahn & Kern, 2005; Mähler *et al*., 2017; Wollenberg Valero, 2020). For each of the centrality statistics, we calculated the mean value for all candidate genes (N = 129) (Data S8). We applied a bootstrap approach by randomly sampling 129 *A. thaliana* genes 1,000 times and calculating the mean and 95% CI of each statistic. To further explore connectivity in gene networks, we used the number of gene-gene interactions in gene networks extracted from STRING v.12 database of protein-protein association networks (Szklarczyk *et al*., 2015). Besides gene co-expression, the interactions are evaluated based on gene co-occurrence, databases, experiments, gene fusion, neighborhood, and text mining in STRING database. We required a minimum interaction score (indicating confidence for an interaction) of medium confidence (0.4). First, we calculated the mean number of interactions for all candidate genes (N = 129) (Data S8). We applied a bootstrap approach by 1000x randomly sampling 129 of *A. thaliana* genes and calculating the mean number of interactions and 95% CI for each set of genes. Second, we grouped genes according to their associations with exchangeable cations identified by GDM analysis (see Figure S4 for the identification of two groups according to their similarity). For the genes associated with macronutrients (N = 53) and genes associated with metals (N = 76) we applied the same approach as for all candidate genes. Lastly, we tested the differences in the number of gene-gene interactions between these two groups by Wilcoxon rank-sum test using the ‘wilcox.test’ function in R package ‘stats v.3.6.2’. Furthermore, we investigated how broadly pleiotropic genes are expressed by determining their tissue specificity (τ specificity index, (Yanai *et al*., 2005)). For pleiotropic genes we expect a low tissue specificity with small τ values. We adopted dataset from (Nocchi *et al*., 2024; Whiting *et al*., 2024) with calculated τ metric (Yanai *et al*., 2005) quantifying the overall tissue specificity of gene expression in *A. thaliana* (across all 23 tissue types and developmental stages; Data S8). Tissue specificity is negatively correlated with pleiotropy as pleiotropic genes are expected to have a low tissue specificity (Yanai *et al*., 2005). Further, for each of the centrality statistics and τ metric we calculated the mean number for all candidate genes (N = 129). We applied a bootstrap approach by 1000x randomly sampling 129 of *A. thaliana* genes and calculating the mean and 95% CI of each statistic for each set of genes.

### Genetic diversity of populations

We evaluated genetic diversity by comparing observed heterozygosity (H_o_) based on genome-wide exonic SNPs (∼ 390 000 SNPs) with H_o_ based on identified candidate SNPs (814 SNPs). We calculated H_o_ based on genome-wide and candidate SNP types for each of the 43 populations using the ‘popStats’ function with genotype likelihood type in vcflib v.1.0.1.1 (Garrison *et al*., 2022). We tested if SNP type affects H_o_ using ‘lm’ (linear model) function from the R package ‘stats v.3.6.2’ and the ‘emmeans’ function (estimated marginal means) from the R package ‘emmeans v.1.10.3’ (Russell, 2024). Further, we inspected the differences in H_o_ based on candidate SNPs among populations assigned to different soil environments (identified in Figure 1d). To do so, we ran a pairwise Wilcoxon rank sum test with correction for multiple testing (Benjamini & Hochberg) using the ‘pairwise.wilcox.test’ function from the R package ‘stats v.3.6.2’. Furthermore, we explored the genetic differentiation among populations based on candidate SNPs in relation to soil heterogeneity across the landscape, following approach in (Moreno-Letelier & Barraclough, 2015; Sakaguchi *et al*., 2017). To test for isolation by environment (IBE), we performed a partial Mantel test with 999 permutations using the ‘mantel.partial’ function from the R package ‘vegan v.2.6.4’ (Oksanen, 2022). This test examines the correlation between genetic (based on candidate SNPs) and environmental distance while controlling for geographic distance. The genetic distances were calculated from population allele frequencies (814 SNPs) using Nei’s genetic distance via ‘nei.dist’ function from the R package ‘poppr v.2.9.6’. We used soil variables for the estimation of environmental distances between populations. Already calculated principal scores 1 (PC1) and 2 (PC2) from PCA based on soil variables were used to compute the Euclidean distances between populations using ‘dist’ function from the R package ‘stats v.3.6.2’. Similarly to pRDA, we accounted for the spatial autocorrelation of populations by including distances computed based on principal component 1 (PC1) scores extracted from genome-wide PCA.

**Figure 1:**
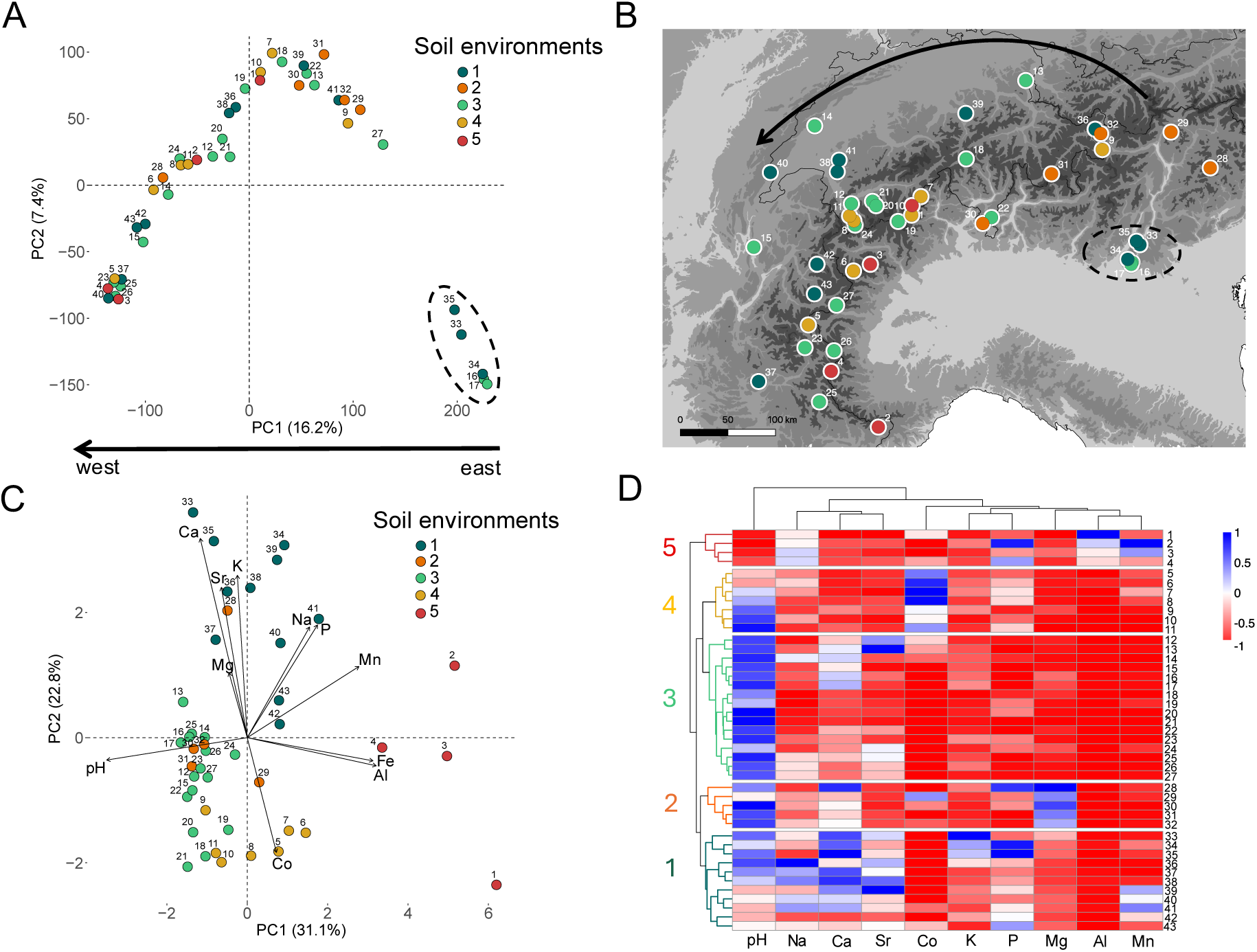
Genetic structure and drivers of soil adaptation in Dianthus sylvestris. A) PCA of 43 populations based on whole genome population allele frequencies at ∼10 millions SNPs shows clustering according to geography (as previously shown in (Luqman et al., 2023)) but not soil type. B) Locations of 43 populations, coloured according to the soil environment, across the Alps. C) PCA biplot of populations based on exchangeable cations. D) Heatmap of standardized exchangeable cations and pH (from 0 to 1) of 43 populations (in rows) clustered based on their similarity. Note: Dashed ellipses indicate area of the glacial refugium and arrows show direction of postglacial range expansion. Populations are coloured according to the soil environments identified in 1d, based on the populations’ similarity in exchangeable cations and pH.

## Results

### Genetic structure and drivers of selection

During range expansion from a calcareous refugial area, alpine populations of *D. sylvestris* colonized a heterogenous mosaic of soil conditions varying in pH and exchangeable cations of macro- and micronutrients and potentially toxic elements such as Al (Figure 1, Figure S4). We clustered the populations into five soil environments according to the similarity based on soil variables (exchangeable cations and pH; Figure 1d). PCA analysis revealed that based on whole genome allele frequencies populations clustered predominantly based on geography-spatial proximity (consistent with (Luqman *et al*., 2023)) and not soil environments, pointing to repeated colonisations of these environments (Figure 1a). We found populations from different soil environments scattered across the Alps, with soil environments 1 and 3 representing refugial soil conditions present from the refugium towards the west (Figure 1a, b). Soil environments 2, 4, and 5 likely represent new conditions that populations encountered during the range expansion. PCA informed by soil variables showed that PC1 explained 31.1% of the variation, mainly reflecting population differences in exchangeable Al, Fe, and pH (Figure 1c). PC2 explained 22.8% of the variation, represented mainly by population differences in exchangeable Ca, Co, K, Mg, and Sr (Figure 1c). The relatively high negative correlations of pH with Al, Fe, and Mn (R² = −0.69, R² = −0.70, and R² = −0.78, respectively; Figure S5) highlight the role of pH in determining the availability of these elements, which may be toxic to plants at high levels in soils with low pH. With the exceptions of strong associations mentioned above and the highest correlation between Al and Fe (R² = 0.87), overall, the correlations among exchangeable cations and pH indicated moderate to weak associations (R² < |0.5|). Therefore, it allows us to explore the effects of specific elements.

To decompose the contributions of soil variables, climate, and population genetic structure (which mirrors geography in the Alps) to genetic variation, we applied partial RDA (pRDA). In the full model, soil variables and population genetic structure significantly explained 36% of the total genetic variation (*p* < 0.001; Table S1a). In the soil model, the effect of soil variables was significant with explaining 20% of total variance (*p* = 0.04; Table S1a) in the data even when accounting for population genetic structure as covariate. Further, the population genetic structure model with soil variables as covariates accounted for 10% of total variance (*p* < 0.001; Table S1a). In the climate model, climate variables explained a similar proportion of total variance like soil variables in the soil model (19% and 18% respectively, Table S1b). Overall, climate variables, soil variables, and population genetic structure shape genomic variation in *D. sylvestris* populations across the Alps.

### Polygenic basis of soil adaptation with varying magnitude of allelic turnover along soil elemental gradients

To identify genetic variants that likely contribute to soil adaptation, we searched for the strongest associations between population allele frequencies with soil variables while accounting for the effect of population structure. Performing univariate environmental association analyses (LFMM2) with each of the 11 soil variables (Al, Ca, Co, Fe, K, Mg, Mn, Na, P, Sr, and pH), we identified 4823 SNPs significantly associated with at least with one soil variable (Table 1, Figure S6, Data S2, and Data S3). The highest overlap in candidate SNPs was observed between Al-associated and Fe-associated SNPs with 368 overlapping SNPs (Figure S6). We excluded Fe-associated SNPs because they were driven only by two populations (Data S1). Thus, we further analysed 3886 soil-associated SNPs occurring in 3158 *D. sylvestris* genes (all filtration steps are described in the Methods; Table S2, Data S3). Overall, most SNPs were associated with Al (1380 SNPs), K (1093 SNPs), Mg (536 SNPs), and Mn (556 SNPs) (Table S2).

**Table 1:**
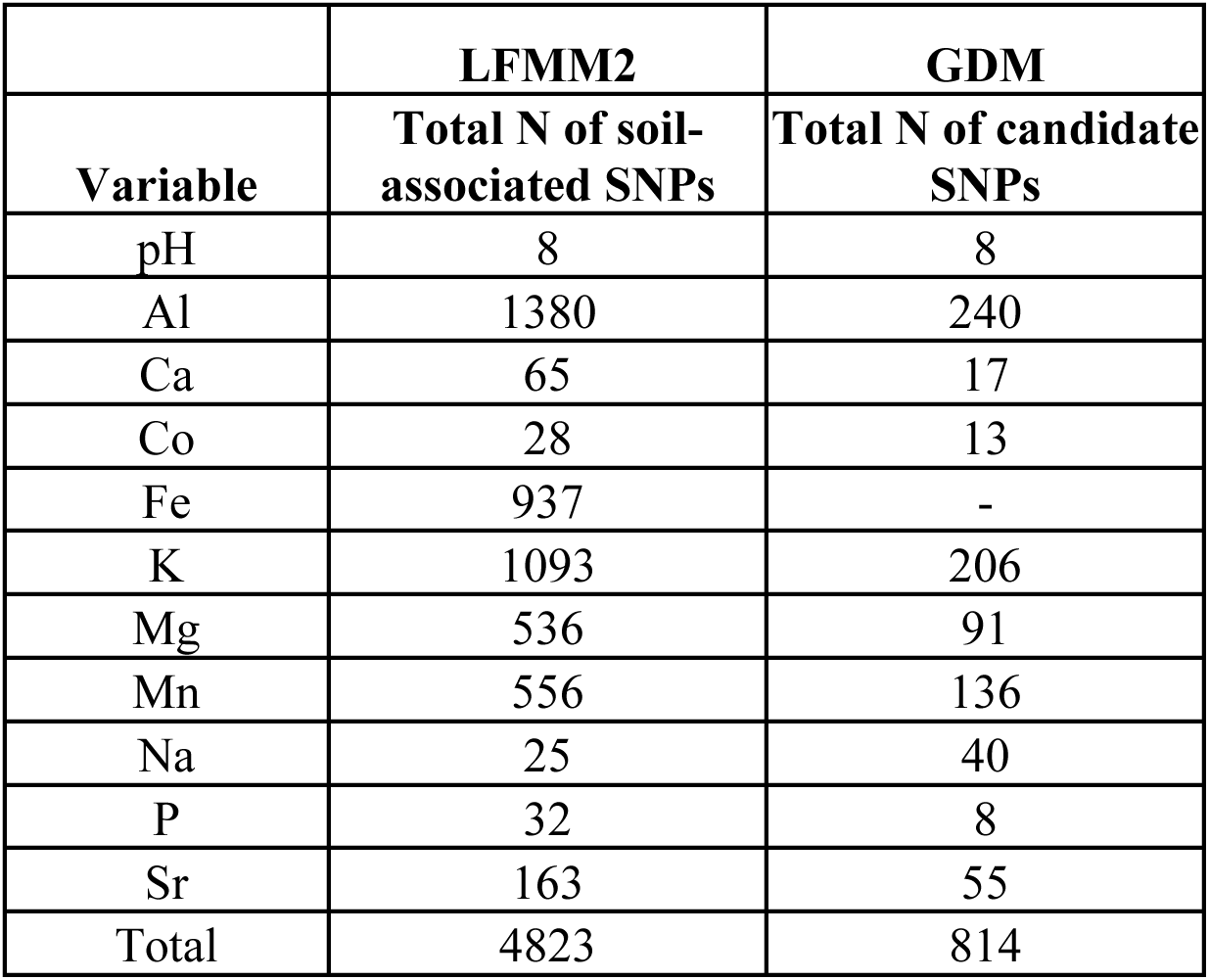
Numbers of identified SNPs associated significantly with soil variables (Al, Ca, Co, Fe, K, Mg, Mn, Na, P, Sr, and pH) in LFMM2 and total numbers of candidate SNPs identified by GDM. Filtration steps are described in the Methods. Note: we excluded Fe-associated SNPs, which resulted in 3886 soil associated SNPs.

|  | LFMM2 | GDM |
| --- | --- | --- |
| Variable | Total N of soil-associated SNPs | Total N of candidate SNPs |
| pH | 8 | 8 |
| Al | 1380 | 240 |
| Ca | 65 | 17 |
| Co | 28 | 13 |
| Fe | 937 | - |
| K | 1093 | 206 |
| Mg | 536 | 91 |
| Mn | 556 | 136 |
| Na | 25 | 40 |
| P | 32 | 8 |
| Sr | 163 | 55 |
| Total | 4823 | 814 |

We investigated the allelic turnover and variation in the rate of change in allele frequencies along soil elemental gradients in 3886 soil-associated SNPs identified by LFMM2 using generalized dissimilarity models (GDM). We included 10 soil variables and geographic distance as predictors in GDM. We then applied filtrations resulting in a dataset of 814 candidate SNPs (see Methods for details; Table 1 and Figure S3). Overall, we kept SNPs with the strongest associations with soil variables identified with the highest variance explained by GDM. Associations with specific soil variables were inferred based on the highest allelic turnover from GDM (Data S4). We presented partial allelic turnovers in candidate SNPs for each soil variable (Figure 2). Specifically, we observed the highest magnitude of allelic turnover, indicated by the height of allelic turnover functions, for K-associated SNPs (mean = 0.97 ± 0.51; 206 SNPs), Al-associated SNPs (mean = 0.73 ± 0.31; 240 SNPs), and Mg-associated SNPs (mean = 0.67 ± 0.36; 91 SNPs). In comparison, the lowest magnitude of allelic turnover was observed for Co-associated SNPs (mean = 0.42 ± 0.18; 13 SNPs), P-associated SNPs (mean = 0.36 ± 0.15; 8 SNPs), and pH-associated SNPs (mean = 0.3 ± 0.13; 8 SNPs) (see Table S3 for complete overview). Moreover, from the shape of the turnover function, we observed a variable rate of change in allele frequencies in candidate SNPs along soil elemental gradients (Figure 2).

**Figure 2:**
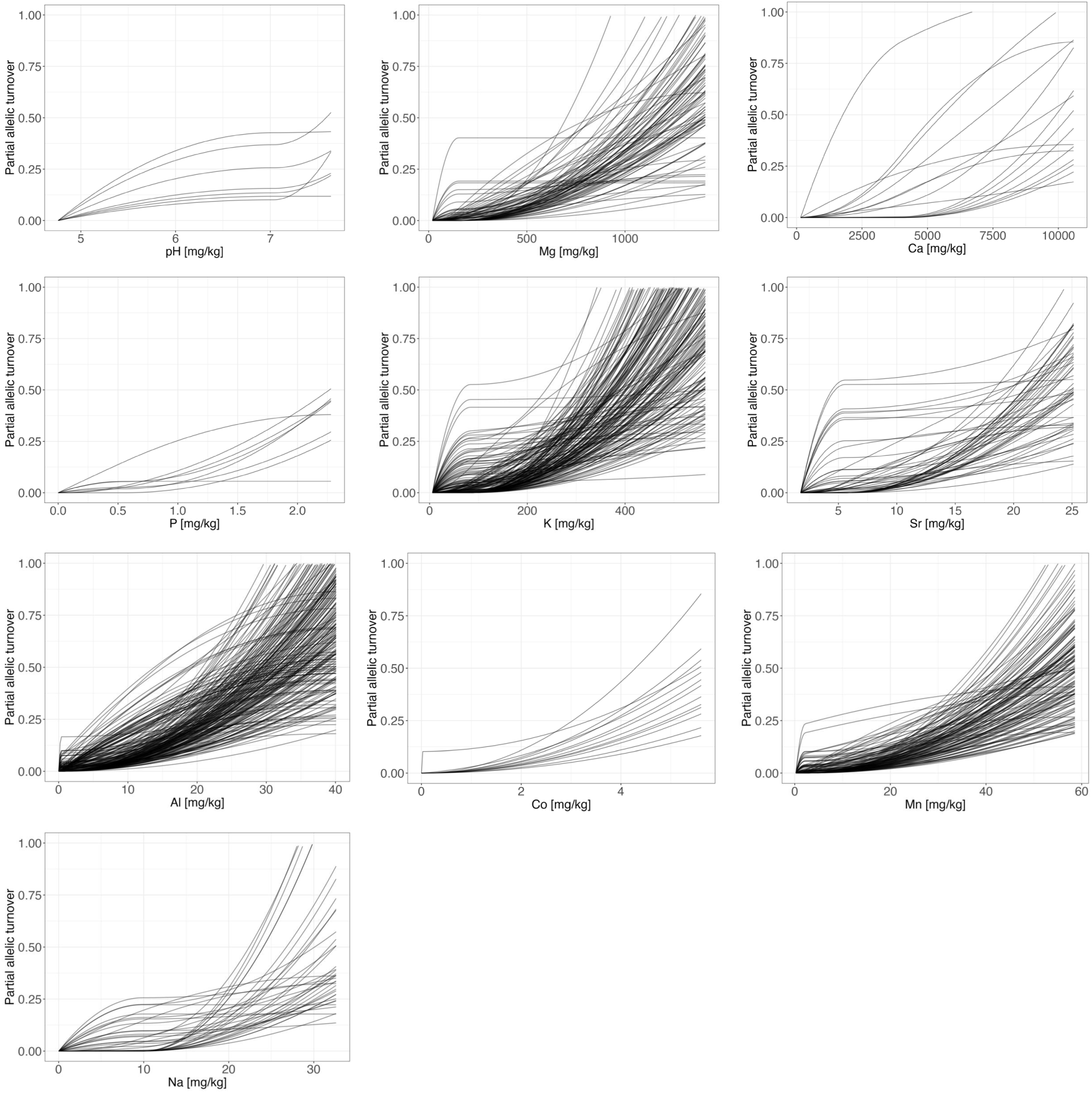
Partial allelic turnover in candidate SNPs along the soil elemental gradients. Allelic turnover functions (GDM fitted I-splines) for candidate SNPs (black lines) that showed the highest partial allelic turnover for particular soil variable. Association of these candidate SNPs with soil variables was inferred based on the relative importance of soil variables in GDM.

### Adaptive genetic composition shaped by selection imposed by soil variables

We explored the relationship between polygenic scores and exchangeable cations, which led to the highest allelic turnovers. Polygenic scores (PS) were shaped by selection in response to K, Mg, and Al exchangeable cations (Figure 3). We observed a strong population differentiation in all three cases and the highest PS based on K-associated SNPs and Al-associated SNPs (Figure 3a, c). The steep slope of the curve indicates selection along exchangeable K and Al (Figure 3a, c). In contrast, the population differentiation was relatively weak based on Mg-associated SNPs, specifically at low Mg levels reflected by the flat curve (Figure 3b). Overall, polygenic scores illustrated how their joint effect reflects variation among populations in exchangeable cations.

**Figure 3:**
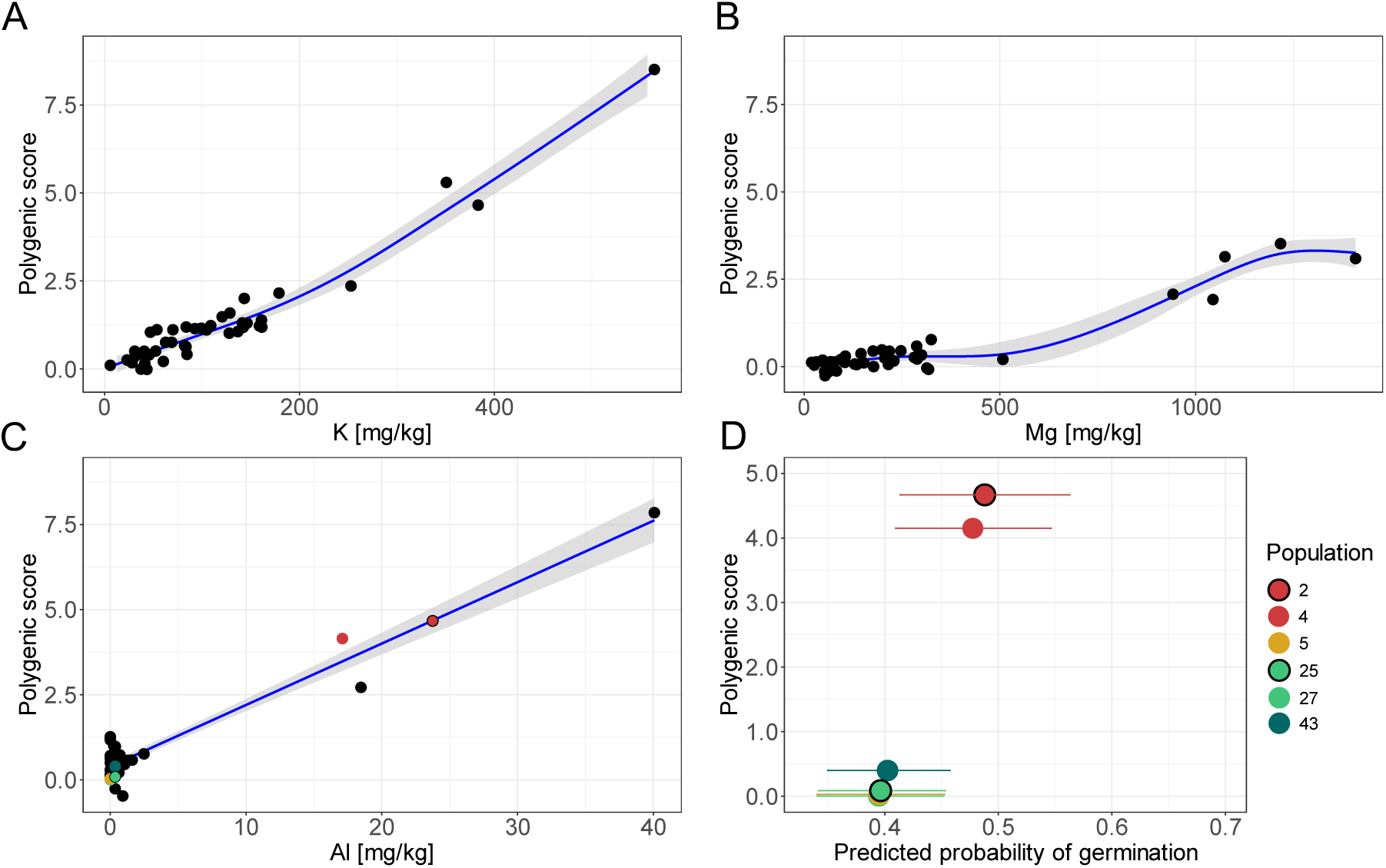
Polygenic scores of 43 populations shaped by selection in response to exchangeable K, Mg, and Al. A) Variation in polygenic scores based on K-associated SNPs (N = 206) along K gradient. B) Variation in polygenic scores based on Mg-associated SNPs (N = 91) along Mg gradient. C) Variation in polygenic scores based on Al-associated SNPs (N = 240) along Al gradient; populations included in the germination experiment are highlighted in colours according to their soil environment. D) Predicted results of binomial generalized linear mixed effect regression model on the probability of seed germination in soils with high Al levels (17 mg/kg and 24 mg/kg). We tested how the adaptive genetic composition (expressed as polygenic scores) of six populations affected germination in high Al level soils (polygenic scores from Figure 3c; in total 768 seeds from six populations; 384 seeds/soil treatment) while accounting for two Al soil treatments and Petri dish as a random effect. Populations are coloured according to soil environments identified in Figure 1d. The bars indicate 95% CI. Source values are presented in Table S4. Note: We calculated polygenic scores with standardized effect size (extracted from LFMM2) allowing comparison among different soil variables.

We observed strong differentiation among populations based on polygenic scores associated with exchangeable Al, specifically with strong selection at high Al levels (Figure 3c). As Al can be toxic to plants in acidic soils (pH < 5), we conducted a germination experiment with six populations (in total 768 seeds) to test how adaptive genetic composition of populations affects seed germination in soils with high Al levels (17 mg/kg and 24 mg/kg). The overall germination rate was 43% in these two soil treatments (44% germination rate in 17 mg/kg and 41% germination rate in 24 mg/kg treatment). We found an association between population polygenic scores and germination using a binomial generalized linear mixed effect regression model, accounting for soil treatment and Petri dish as a random factor (estimate = 0.08138, SE = 0.03627, *p* = 0.02485). Further, for each of these six populations based on the output from the tested model, we predicted the probability of seed germination in high Al levels. Overall, we found that populations with the highest polygenic scores, i.e. originally from the soils with high Al levels (Population 2 and 4), had higher germination rates in soils with high Al levels than populations originally from low Al soils (Figure 3d, Table S4, Data S5).

### Functions of candidate genes in soil adaptation

In total 814 candidate SNPs were aligned to 770 *D. sylvestris* genes. To further inspect putative functions and relevance of the candidate genes for soil adaptation, we identified 475 *Arabidopsis thaliana* orthologs by reciprocal best blast hit between protein sequences of *D. sylvestris* and *A. thaliana.* Out of these, we applied a strict threshold of at least 70% identity in the alignment, which resulted in 129 candidate genes (Data S6). Gene ontology analysis revealed significantly enriched (p < 0.05) biological processes (BP) and molecular functions (MF) (in total 80 BP and 28 MF, overview in Data S7). For further interpretation, we explored a subset of the significantly enriched GO terms relevant for soil adaptation (see Methods for criteria) including ion transport and homeostasis (response to copper ion, manganese ion transmembrane transport, calcium ion transmembrane transport, organic anion transport, copper ion binding, iron-sulfur cluster binding, metal ion homeostasis, intracellular monoatomic ion homeostasis, monoatomic cation homeostasis, and inorganic ion homeostasis), drought response (stomatal complex development), and root growth (cell tip growth) (Figure S7).

We explored the list of candidate genes (N = 129) and refined the selection of the genes for further interpretations (Table 2) – we focused on genes annotated to multiple enriched GO terms (from a subset of GO terms relevant for soil adaptation; Data S7, Figure S7) and the gene *NRT2.4* with the highest effect size (in absolute values; extracted from the LFMM2, Data S6). Additionally, we checked if the selected ion transporter genes are expressed in the root tissues ((Papatheodorou *et al*., 2018); Expression Atlas). Based on these criteria, the refined list includes ion transporter genes, such as *SKOR* (AT3G02850), a member of Shaker family potassium ion (K^+^) channel and *HMA5* (AT1G63440), which is involved in Cu detoxification (del Pozo *et al*., 2010). Further, *NRT2.4* (AT5G60770) as a high-affinity nitrate transporter regulated in *A. thaliana* by nitrogen availability, with its expression induced by nitrogen deficiency (Berardini *et al*., 2004; Zhuo *et al*., 2024). In this study, variation in *NRT2.4* was associated with another macronutrient – Ca. We found a correlation between population allele frequencies at this candidate gene (specifically SNP_scaffold426_size137742_1_125730_35650, which had the highest effect size among candidates) and exchangeable Ca in soil (cubic regression, R^2^ = 0.51; p < 0.001). This suggests different responses of populations to varying exchangeable Ca in soil (Figure 4). The highest rate of change in allele frequencies, indicated by the steep slope of the spline, was observed at low soil Ca levels, with the highest allele frequencies found in populations originating from low soil Ca levels (Figure 4). Another ion transporter, *CAX5* (AT1G55730), a member of low affinity calcium antiporter CAX2 family, is up-regulated in response to high Mn, Ca, and Na stress (Edmond *et al*., 2009). We also identified *PML5* (AT1G68650), Mn/Ca transporter, and *MTP11* (AT2G39450), which encodes a protein in *A. thaliana* involved in Mn transport and tolerance via vesicular trafficking and exocytosis (Peiter *et al*., 2007). Furthermore, we found genes involved in stomatal complex development, *EPF1* (AT2G20875) and *TUBG2* (AT5G05620), a process putatively related to the response to drought. Lastly, we found gene *SKU5* (AT4G12420), which is involved in cell tip growth and is presumably linked to root growth.

**Table 2:** Overview of selected candidate genes for soil adaptation. Gene names and descriptions were extracted from the Araport11 annotation (available in TAIR database www.arabidopsis.org). Effect sizes were extracted from LFMM2. Population allele frequency is the mean allele frequency across all 43 populations with the minimum and maximum allele frequency in the brackets. Soil variable association was inferred from the GDM. Expression of particular candidate genes in root tip and in root tissue (in TPM - transcripts per kilobase million) were extracted from (Papatheodorou et al., 2018) (available in Expression Atlas www.ebi.ac.uk).

| Gene ID | Gene name | Description | Effect size | Population allele frequency | Soil variable association | Expression in root tip [TPM] | Expression in root [TPM] |
| --- | --- | --- | --- | --- | --- | --- | --- |
| AT1G55730 | *CAX5* | *CAX5* is a member of low affinity calcium antiporter *CAX2* family. | 0.04 | (0)0.06(0.29) | Mn | 26 | 27 |
| AT1G63440 | *HMA5* | P-type ATPase *HMA5* is involved in Cu detoxification. | 0.05 | (0)0.07(0.36) | Na | 25 | 77 |
| AT1G68650 | *PML5* | Member of the UPF0016 family of membrane proteins, belongs to the conserved group of Mn/Ca transporters. | 0.10 | (0)0.31(0.88) | Mg | 25 | 24 |
| AT2G20875 | *EPF1* | *EPF1* encodes a secretory peptide *EPF1* involved in stomatal development. | 0.09 | (0.21)0.49(0.85) | Mg | 0.1 | 0 |
| AT2G39450 | *MTP11* | *MTP11* encodes a Golgi-localized manganese transporter that is involved in Mn tolerance. | 0.06 | (0)0.19(0.57) | Mn | 71 | 101 |
| AT3G02850 | *SKOR* | *SKOR* is a member of Shaker family potassium ion (K<sup>+</sup>) channel. It mediates the delivery of K<sup>+</sup> from stelar cells to the xylem in the roots towards the shoot. | 0.05 | (0)0.14(0.38) | Al | 42 | 4 |
| AT4G12420 | *SKU5* | *SKU5* encodes a protein of unknown function involved in directed root tip growth. | -0.06 | (0.38)0.65(0.88) | Al | 165 | 192 |
| AT5G05620 | *TUBG2* | Paralog of *TUBG1* is required for centrosomal and noncentrosomal microtubule nucleation. It is involved in specification of cell identity, such as stomatal patterning. | 0.07 | (0)0.14(0.36) | K | 23 | 34 |
| AT5G60770 | *NRT2.4* | *NRT2.4* is a member of high affinity nitrate transporter family. | -0.15 | (0)0.22(0.85) | Ca | 0.5 | 0.5 |

**Figure 4:**
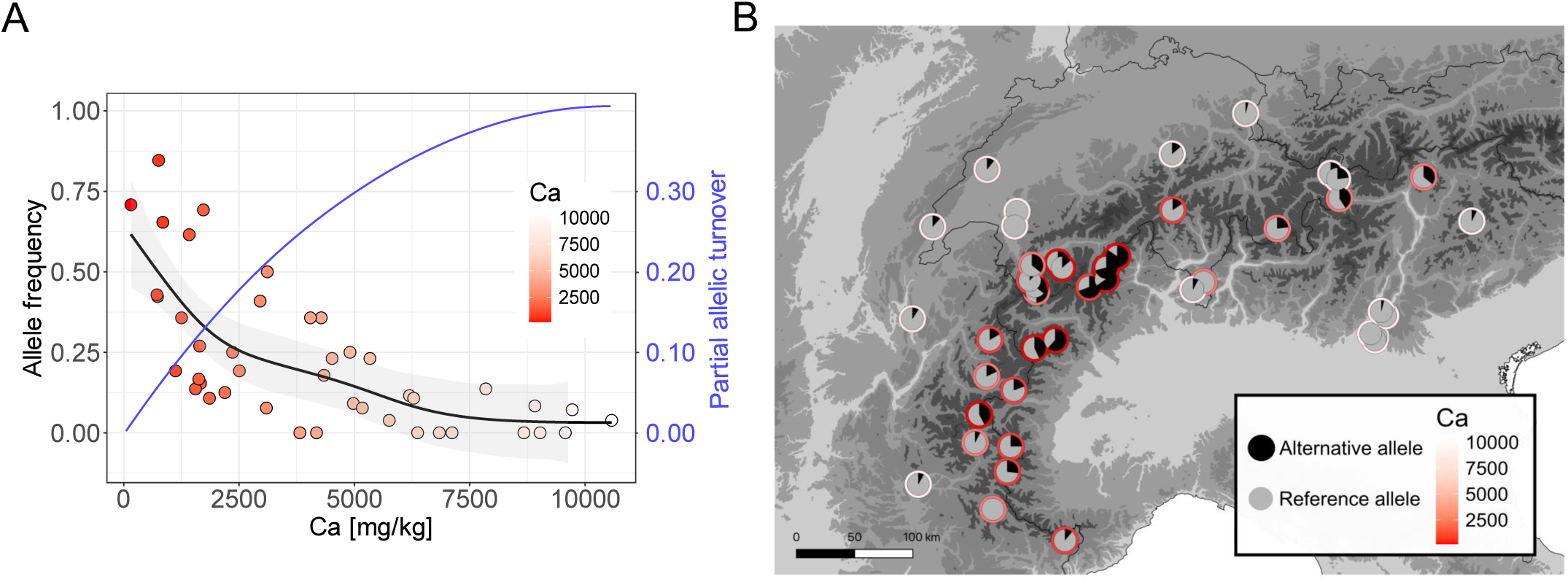
Genetic variation in NRT2.4 candidate gene. A) Correlation of population allele frequencies (left y-axis) of candidate SNP (scaffold426_size137742_1_125730_pos35650) with exchangeable Ca (non-linear model with cubic regression, R^2^ = 0.51; p < 0.001); partial allelic turnover of this candidate SNP identified by GDM (right y-axis in blue). B) Variation in allele frequencies of the candidate SNP in 43 populations along the range expansion. Note: red-white colour gradient denotes exchangeable Ca in each population and grey-black colour shows the frequency of reference/alternative allele.

### Pleiotropy facilitates polygenic adaptation to novel selective pressures

We further asked whether candidate genes are of higher pleiotropy than randomly sampled sets of genes. First, we explored the number of gene-gene interactions finding that candidate genes have significantly more interactions (mean: 265) than the other *A. thaliana* genes (mean: 125, 95% CI: 99-154; 1000 permutations, *p* < 0.001; Figure 5, Table S5). Interestingly, candidate genes associated with metals (N = 76, mean: 287) appeared to be more interconnected than genes associated with macronutrients (N = 53, mean: 233; Figure 5, Table S5), although the difference in the number of gene-gene interactions was not significant (Wilcoxon rank-sum test, W = 1654, *p* = 0.1182). Second, we examined pleiotropy by various centrality statistics to test if candidate genes are more central in gene networks than randomly sampled genes. We found significantly higher node closeness (mean: 9.7 × 10^−6^), node degree (mean: 697), and node strength (mean: 2374) of candidate genes within gene co-expression networks than other *A. thaliana* genes (mean node closeness: 9.3 × 10^−6^, 95% CI: 9.2 × 10^−6^-9.46 × 10^−6^; mean node degree: 473, 95% CI: 426-526; mean node strength: 1559, 95% CI: 1378-1751; in all cases *p* < 0.001; Table 3). The only exception was node betweenness, which was not significantly higher for candidate genes (mean: 26566) than for the other genes (mean: 24871, 95% CI: 22237-27711; *p* = 0.1229; Table 3). Third, we explored tissue specificity of candidate genes expression. We found significantly lower tissue specificity of candidate genes (τ = 0.62) than other *A. thaliana* genes (τ = 0.76, 95% CI: 0.72-0.79, *p* < 0.001; Table 3), i.e. candidate genes are possibly more broadly expressed within gene co-expression networks implying possibly higher pleiotropy. Overall, candidate genes were more interconnected, with more central positions in gene networks than other genes in co-expression networks, and they had a lower tissue specificity.

**Figure 5:**
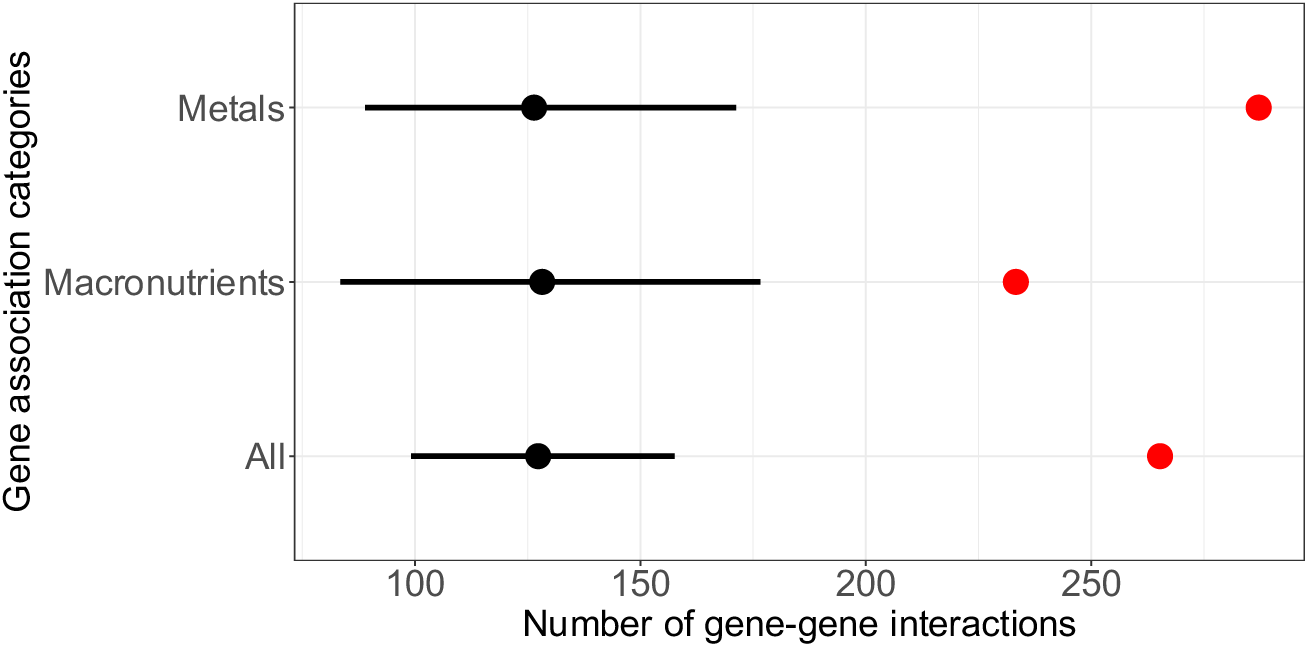
Gene-gene interactions of 129 candidate genes. Genes were classified according to their associations inferred by GDM (Metals: 76 genes, Macronutrients: 53 genes; Data S6). Red dots indicate mean number of candidate gene interactions (inferred from STRING database) within gene networks. Black dots and lines represent mean and 95% quantiale of number of gene interactions among 1000x randomly sampled genes of A. thaliana. Note: Original values are presented in Table S5.

**Table 3:**
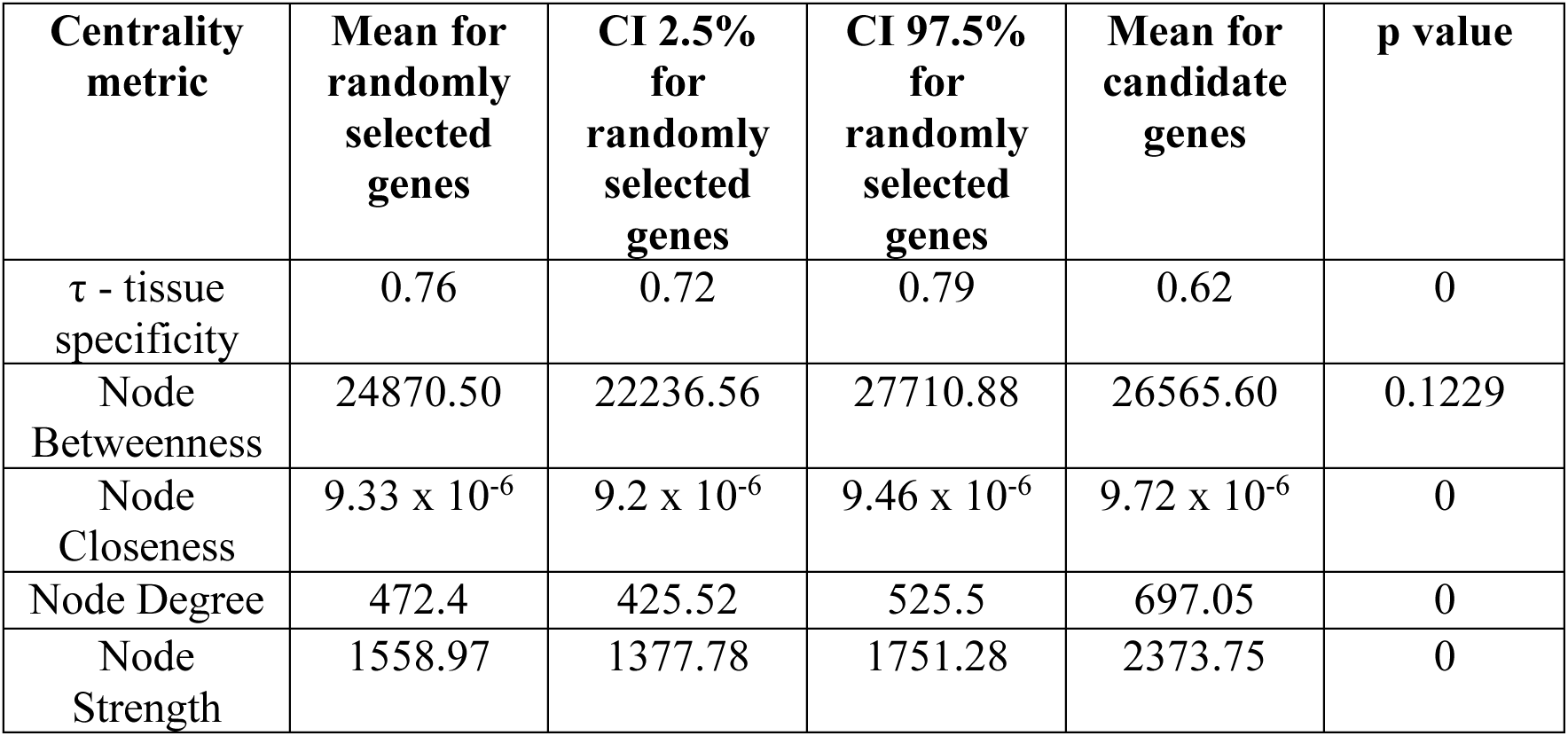
Degree of pleiotropy of candidate genes. Overview of centrality metrics and tissue specificity (τ) used as proxy for pleiotropy. Random set of genes (with the same number as candidate genes) were sampled 1000X for each metric.

| Centrality metric | Mean for randomly selected genes | CI 2.5% for randomly selected genes | CI 97.5% for randomly selected genes | Mean for candidate genes | p value |
| --- | --- | --- | --- | --- | --- |
| $\tau$ - tissue specificity | 0.76 | 0.72 | 0.79 | 0.62 | 0 |
| Node Betweenness | 24870.50 | 22236.56 | 27710.88 | 26565.60 | 0.1229 |
| Node Closeness | $9.33 \times 10^{-6}$ | $9.2 \times 10^{-6}$ | $9.46 \times 10^{-6}$ | $9.72 \times 10^{-6}$ | 0 |
| Node Degree | 472.4 | 425.52 | 525.5 | 697.05 | 0 |
| Node Strength | 1558.97 | 1377.78 | 1751.28 | 2373.75 | 0 |

### Soil heterogeneity helps to maintain adaptive genetic variation in populations

We compared populations observed heterozygosity (H_o_) based on genome-wide exonic SNPs and candidate SNPs. Despite the observed general trend in decreasing genetic diversity along range expansion (Figure S8), we found overall higher population-level H_o_ calculated based on candidate SNPs than based on genome-wide exonic SNPs (linear model; estimate = 0.081, SE = 0.0258, *p* = 0.0023; Figure 6a). Further, we explored differences in H_o_ based on candidate SNPs among populations assigned to different soil environments (Figure 6b, c). We observed the highest mean H_o_ (H_o_ = 0.21; Table S6) for soil environment 5, which includes populations originally from acidic soils according to low soil pH values. In contrast, soil environment 3, which includes the majority of populations originating from the alkaline soils according to high soil pH values, showed the lowest mean H_o_ (H_o_ = 0.17; Table S6). We found significant differences in H_o_ based on candidate SNPs between soil environment 1 - 3 (pairwise Wilcoxon rank sum test, *p* = 0.028), soil environment 3 - 5 (pairwise Wilcoxon rank sum test, *p* = 0.017), and soil type 4 - 5 (pairwise Wilcoxon rank sum test, *p* = 0.04; summary of *p*-values for all pairwise comparisons is presented in Table S7). Furthermore, given these differences among soil environments, we tested if the genetic distance based on candidate SNPs correlates with environmental distance based on soil variables. We revealed a pattern of isolation by environment (IBE) – a significant positive correlation between genetic and environmental distances (partial Mantel test R = 0.75, *p* < 0.001; Figure 6d). This highlights the effects of soil heterogeneity in the landscape in shaping the populations’ adaptive genetic variation in the space. Overall, our findings suggest that spatially varying selection in a heterogeneous soil landscape contributes to the maintenance of adaptive genetic variation in populations.

**Figure 6:**
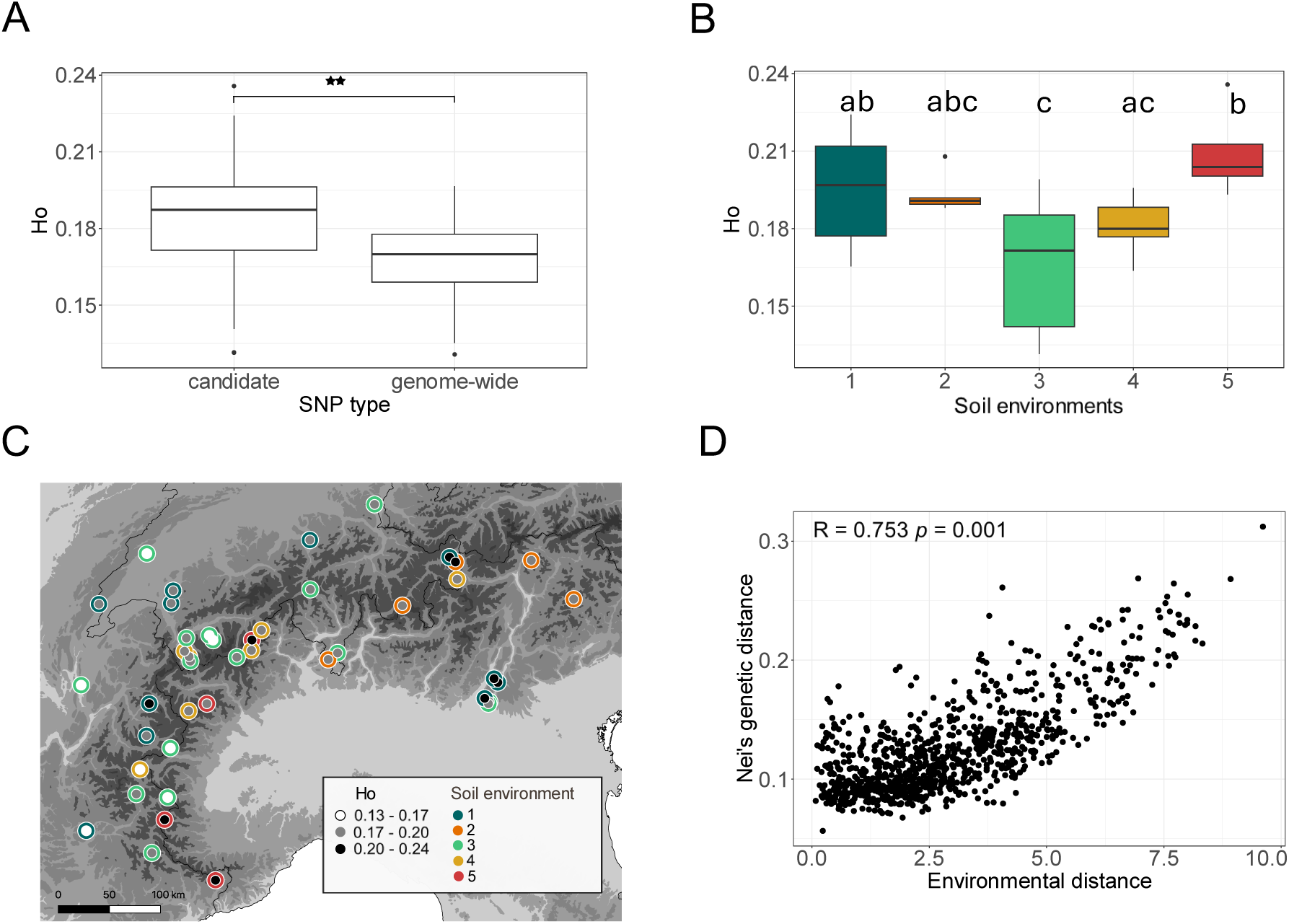
Genetic diversity of populations in heterogeneous mosaic of soil conditions. A) Difference in populations observed heterozygosity (H_o_) based on candidate SNPs and genome-wide exonic SNPs (asterisks denote the significance of the SNP type effect, ∗∗ p = 0.0023). B) Differences in H_o_ based on candidate SNPs among populations grouped according to different soil environments (identified in Figure 1d). Different letters indicate significant differences for pairwise comparisons (p < 0.05), an overview of corrected p-values for multiple testing is in Table S7. C) Locations of populations along the range expansion with presented H_o_ based on candidate SNPs. D) Isolation by the environment – partial Mantel test on environmental (soil) and genetic distances between populations.

## Discussion

Here, we studied the polygenic basis of adaptation to contrasting soil conditions in *D. sylvestris* leveraging low-coverage genome-wide data from 43 populations across the Alps. We found the highest allelic turnover in candidate SNPs associated with K, Mg, and Al. In potentially toxic soils with high Al levels, we found that populations with high polygenic scores based on Al-associated alleles had a higher germination rate than populations with low polygenic scores. Candidate genes for soil adaptation showed higher pleiotropy with more central positions in the gene networks than randomly sampled genes, pointing to an important role of pleiotropy in driving polygenic adaptation to new environmental conditions. Soil heterogeneity likely played an important role in maintaining adaptive genetic variation in populations during the range expansion in the Alps.

### Soil variables as drivers of selection

We show that soil factors play a significant role in shaping genetic variation during range expansion of *D. sylvestris*. Interestingly, climate and soil variables explain a similar proportion of variation in the genetic data (Table S1b), highlighting their joint influence on selection during range expansion. This is consistent with previous studies (Luqman *et al*., 2023; Paalsson *et al*., 2024), which showed that range expansion in *D. sylvestris* was linked to the colonization of heterogeneous landscapes and adaptation to new environments such as warm habitats in alpine valleys. While adaptation to climatic conditions in mountain regions has been extensively studied e.g., (de Villemereuil *et al*., 2018; Rellstab *et al*., 2020; Wos *et al*., 2022) soil adaptation has received less attention, even though soil conditions can be highly heterogeneous and influence vegetation composition (Takahashi & Murayama, 2014). We demonstrate that soil variables drive selection on many alleles (Figure 2). We explored allelic turnover along soil elemental gradients, finding high differences in allelic turnovers at high levels of nutrients and metals. Therefore, they are important drivers of genomic variation, suggesting strong responses that likely influence adaptation in populations with high levels metals or nutrients. However, different candidate SNPs for the same elements showed the highest rate of change in allele frequencies at low levels (Figure 2). Importantly, the high variation in responses along soil elemental gradients for each soil variable suggests that different sets of candidate SNPs may contribute to adaptation at low and at high soil metal and nutrient soil levels.

Overall, we found the highest turnover rates in SNPs associated with K, Mg, and Al (Table S3). Potassium, the most abundant cation required for plant growth, plays a crucial role in processes such as plant signalling and stomatal regulation (Wang & Wu, 2013; Mostofa *et al*., 2022). K homeostasis also influences other nutrients and is critical for plant responses to abiotic stresses, including drought and metal toxicity (Naciri *et al*., 2021; Mostofa *et al*., 2022; Yan *et al*., 2023). Specifically, we observed the positive effect sizes for most K-associated SNPs with Ca, Mg, and P, highlighting their synergistic interactions. Further, 55% of SNPs associated with both Al and K had positive effect sizes, while 41% showed different directions, suggesting both synergistic and antagonistic effects (Figure S6, Data S3). Magnesium is essential for plant development, photosynthesis, and nutrient metabolism (Ahmed *et al*., 2023). It plays a crucial role in the uptake and transport of nutrients like P and regulates Ca and K ions transport (Sigel & Sigel, 1998). This likely explains the high allelic turnover observed in K- and Mg-associated SNPs (Tables S3). In fact, 70% of SNPs associated with both K and Mg exhibited positive effect sizes pointing to synergistic effects (Figure S6, Data S3). Aluminium is particularly challenging for plants in acidic soils (pH < 5), inhibits root growth, induces oxidative stress, and disrupts nutrient uptake due to increased exchangeable Al (Clarkson, 1965; Ryan *et al*., 2011; Gould *et al*., 2014). Consistent with this, we found the strongest selection at high Al levels (Figure 3c).

### Soil variables impose changes in the adaptive genetic composition of populations

Theory predicts that polygenic traits are expressed through the combined effect of many alleles. Here, we used polygenic scores to summarize the adaptive genetic composition of populations based on SNPs associated with soil elements that exhibited the highest allelic turnover. We found a strong association of polygenic scores with exchangeable K, Mg, and Al (Figure 3). It suggests that these elements act as selective pressures, likely driving polygenic soil adaptation in *D. sylvestris* populations. The polygenic score approach has been previously applied mainly to study the cumulative contribution of candidate genes associated with specific climatic variables (Beer *et al*., 2022; Folkertsma *et al*., 2024; Van Deurs *et al*., 2025). While many studies have identified genotype-by-environment associations, the experimental testing and prediction of their functional effects remain rare, with some exceptions e.g. (Lasky *et al*., 2015; Troth *et al*., 2018). Here, following other studies on soil adaptation, which found that high Al levels negatively impacted root growth in *Anthoxanthum odoratum* (Gould *et al*., 2014) and *Sorghum bicolor* (Lasky *et al*., 2015), we conducted a germination experiment with high soil Al levels focusing on the germination phase, which is known to be highly sensitive to soil chemical composition (Guggisberg *et al*., 2018). In line with expectations, populations originating from soils with high Al levels exhibited higher germination rates in high Al soils than populations from low Al soils (Figure 3d). This supports the idea that populations from high Al soils are likely to carry more Al-associated alleles than populations with lower polygenic scores, which originate from low Al soils.

### Role of pleiotropic genes in adaptation

We showed that candidate genes are more interconnected with more central positions in gene networks and exhibit lower tissue specificity compared to other genes in co-expression networks (Table 3, Figure 5). Therefore, pleiotropy may play a significant role in the regulation of both essential and nonessential metal ions and nutrients. Specifically, genes associated with metals were the most interconnected, indicating a coordinated response to metal stress, particularly Al (Table S5). As soils with high Al levels represent new environmental conditions for *D. sylvestris*, our findings are congruent with simulation in (Hamala *et al*., 2020) showing that pleiotropy is beneficial when populations are far from their previous optimum, i.e. facing novel selective pressures. Such cases can occur during invasions, as has been shown in *Ambrosia artemisiifolia* (Hamala *et al*., 2020). Similarly to invasions, during range expansions, populations may also experience novel selective pressures, such as during colonization of freshwater habitats in sticklebacks (Rennison & Peichel, 2022) or cold habitats in *Arabidopsis thaliana* (Des Marais *et al*., 2017). In these species and also in *D. sylvestris*, it can be particularly important to maintain genetic variation with synergistic pleiotropic effects, which facilitates rapid adaptation of populations that repeatedly colonize similar environments. In contrast to the traditional view on pleiotropy (Fisher, 1930; Orr, 2000), our findings of pleiotropic candidate genes for adaptation are consistent with recent empirical studies showing the important role of pleiotropy in the adaptive evolution (McGee *et al*., 2016; Frachon *et al*., 2017; Archambeault *et al*., 2020; Hamala *et al*., 2020; Rennison & Peichel, 2022).

### Role of soil heterogeneity in maintaining genetic variation

We found higher observed heterozygosity (H_o_) of populations based on candidate SNPs than based on genome-wide exonic SNPs (Figure 6a). This suggests that selection maintains adaptive genetic variation in soil-associated loci. These adaptive variants may have originated from highly divergent haplotypes already present in the ancestors of *D. sylvestris* (Luqman, 2021). Despite the general trend of decreasing population-level H_o_ along the range expansion (Figure S8), populations from acidic soils (soil environment 5) exhibited the highest H_o_ in candidate SNPs (Figure 6b, Table S6). This indicates that populations, even at the range expansion western edge (Figure 6c), seem to have adapted to novel environmental conditions, as the acidic soils represent new conditions for *D. sylvestris*. Past studies on polygenic adaptation suggest multiple mechanisms for maintaining standing genetic variation in populations (Barton & Turelli, 2004; Yeaman *et al*., 2010). One such mechanism is spatially varying selection in heterogeneous environments (Byers, 2005; Yeaman & Jarvis, 2006; Delph & Kelly, 2014; Yeaman, 2015). Specifically, strong selection despite gene flow is crucial for maintaining polygenic variation in spatially heterogeneous environments (Spichtig & Kawecki, 2004). However, the generality of these findings remains unclear, because even within the same study system, e.g. lodgepole pine, conclusions about the relationship between environmental heterogeneity and genetic variation differ (Yeaman & Jarvis, 2006; Archambeau *et al*., 2023). The environmental heterogeneity, in the case of soil adaptation reflected in differences in soil chemical properties across the landscape, likely contributes to maintaining adaptive genetic variation.

## Conclusions

We explored soil variables as important factors shaping plant adaptation in a heterogenous landscape. Specifically, we examined the polygenic basis of soil adaptation and how novel selective pressures encountered during *Dianthus sylvestris* postglacial range expansion shaped populations’ adaptive genetic composition. We found high differences in allelic turnovers at high levels of nutrients and metals, suggesting that these factors are important drivers of genomic variation. We observed the significant effect of adaptive genetic composition on seed germination in response to toxic high Al-soils. However, further experimental validations will be necessary as a follow-up step. Furthermore, our findings indicate that pleiotropy plays a crucial role in polygenic adaptation to novel selective pressures. Lastly, soil heterogeneity promotes spatial varying selection, which helps to maintain genetic variation presumably needed for the colonization of new environments.

## Acknowledgements

We thank Karim Clivaz and Karsten Rohweder for help with fieldwork and Nicole Helbing and Maike Friedel for assistance with soil processing. We are grateful to Christian Rellstab for initial discussions on environmental association analyses and to the Genetic Diversity Centre at ETH Zurich, in particular Niklaus Zemp for providing IT support. This research was supported by the Swiss National Science Foundation (SNSF) grants 31003A_160123 and 31003A_182675 awarded to A.W. and the Czech Science Foundation grant 23-05469O awarded to V.L. Additional support was provided by the Czech Academy Sciences (long-term research development project RVO 67985939).

## Notes

### Competing Interest Statement

The authors have declared no competing interest.

## References

Ahmed N, Zhang B, Bozdar B, Chachar S, Rai M, Li J, Li Y, Hayat F, Chachar Z, Tu P. 2023. The power of magnesium: unlocking the potential for increased yield, quality, and stress tolerance of horticultural crops. Frontiers in Plant Science 14: 1285512.

Alexa A, Rahnenführer J. 2009. Gene set enrichment analysis with topGO. Bioconductor Improv 27: 1–26.

Alexa A, Rahnenführer J. 2022. topGO: Enrichment Analysis for Gene Ontology. R package version 2.5.

Archambeau J, Benito Garzón M, de Miguel M, Brachi B, Barraquand F, González-Martínez SC. 2023. Reduced within-population quantitative genetic variation is associated with climate harshness in maritime pine. Heredity 131(1): 68–78.

Archambeault SL, Bartschi LR, Merminod AD, Peichel CL. 2020. Adaptation via pleiotropy and linkage: Association mapping reveals a complex genetic architecture within the stickleback Eda locus. Evol Lett 4(4): 282–301.

Barghi N, Hermisson J, Schlötterer C. 2020. Polygenic adaptation: a unifying framework to understand positive selection (vol 18, pg 913, 2020). Nature Reviews Genetics 21(12): 782–782.

Barrett RDH, Schluter D. 2008. Adaptation from standing genetic variation. Trends in Ecology & Evolution 23(1): 38–44.

Barton N, Turelli M. 2004. Effects of genetic drift on variance components under a general model of epistasis. Evolution 58(10): 2111–2132.

Barton NH. 1999. Clines in polygenic traits. Genet Res 74(3): 223–236.

Barton NH, Turelli M. 1989. Evolutionary quantitative genetics: how little do we know? Annual Review of Genetics 23: 337–370.

Bates D, Machler M, Bolker B, Walker S. 2015. Fitting linear mixed-effects models using lme4. Journal of Statistical Software 67: 1–48.

Beer MA, Kane RA, Micheletti SJ, Kozakiewicz CP, Storfer A. 2022. Landscape genomics of the streamside salamander: Implications for species management in the face of environmental change. Evolutionary Applications 15(2): 220–236.

Berardini TZ, Mundodi S, Reiser L, Huala E, Garcia-Hernandez M, Zhang P, Mueller LA, Yoon J, Doyle A, Lander G. 2004. Functional annotation of the Arabidopsis genome using controlled vocabularies. Plant physiology 135(2): 745–755.

Binkley D, Vitousek P 1989. Soil nutrient availability. Plant physiological ecology: field methods and instrumentation: Springer, 75–96.

Bohutínská M, Vlček J, Yair S, Laenen B, Konečná V, Fracassetti M, Slotte T, Kolář F. 2021. Genomic basis of parallel adaptation varies with divergence in Arabidopsis and its relatives. Proceedings of the National Academy of Sciences 118(21): e2022713118.

Bomblies K, Peichel CL. 2022. Genetics of adaptation. Proc Natl Acad Sci U S A 119(30): e2122152119.

Booker TR. 2024. The structure of the environment influences the patterns and genetics of local adaptation. Evolution Letters 8(6): 787–798.

Bridle JR, Vines TH. 2007. Limits to evolution at range margins: when and why does adaptation fail? Trends in Ecology & Evolution 22(3): 140–147.

Byers DL. 2005. Evolution in heterogeneous environments and the potential of maintenance of genetic variation in traits of adaptive significance. Genetica 123: 107–124.

Camacho C, Coulouris G, Avagyan V, Ma N, Papadopoulos J, Bealer K, Madden TL. 2009. BLAST+: architecture and applications. BMC bioinformatics 10: 1–9.

Caniato FF, Guimaraes CT, Hamblin M, Billot C, Rami J-F, Hufnagel B, Kochian LV, Liu J, Garcia AAF, Hash CT. 2011. The relationship between population structure and aluminum tolerance in cultivated sorghum. PLoS One 6(6): e20830.

Capblancq T, Forester BR. 2021. Redundancy analysis: A Swiss Army Knife for landscape genomics. Methods in Ecology and Evolution 12(12): 2298–2309.

Cattell RB. 1966. The scree test for the number of factors. Multivariate behavioral research 1(2): 245–276.

Caye K, Jumentier B, Lepeule J, Francois O. 2019. LFMM 2: Fast and Accurate Inference of Gene-Environment Associations in Genome-Wide Studies. Mol Biol Evol 36(4): 852–860.

Clarkson DT. 1965. Effect of Aluminium and Some Other Trivalent Metal Cations on Cell Division in Root Apices of Allium Cepa. Annals of Botany 29(114): 309.

Crawford K, Whitney K. 2010. Population genetic diversity influences colonization success. Molecular Ecology 19(6): 1253–1263.

Dauphin B, Rellstab C, Schmid M, Zoller S, Karger DN, Brodbeck S, Guillaume F, Gugerli F. 2021. Genomic vulnerability to rapid climate warming in a tree species with a long generation time. Global Change Biology 27(6): 1181–1195.

de Villemereuil P, Mouterde M, Gaggiotti OE, Till-Bottraud I. 2018. Patterns of phenotypic plasticity and local adaptation in the wide elevation range of the alpine plant Arabis alpina. Journal of Ecology 106(5): 1952–1971.

del Pozo T, Cambiazo V, González M. 2010. Gene expression profiling analysis of copper homeostasis in Arabidopsis thaliana. Biochemical and biophysical research communications 393(2): 248–252.

Delph LF, Kelly JK. 2014. On the importance of balancing selection in plants. New Phytologist 201(1): 45–56.

Des Marais DL, Guerrero RF, Lasky JR, Scarpino SV. 2017. Topological features of a gene co-expression network predict patterns of natural diversity in environmental response. Proc Biol Sci 284(1856).

Dormann CF, Elith J, Bacher S, Buchmann C, Carl G, Carré G, Marquéz JRG, Gruber B, Lafourcade B, Leitão PJ. 2013. Collinearity: a review of methods to deal with it and a simulation study evaluating their performance. Ecography 36(1): 27–46.

Dudaniec RY, Yong CJ, Lancaster LT, Svensson EI, Hansson B. 2018. Signatures of local adaptation along environmental gradients in a range-expanding damselfly (Ischnura elegans). Molecular Ecology 27(11): 2576–2593.

Durinck S, Moreau Y, Kasprzyk A, Davis S, De Moor B, Brazma A, Huber W. 2005. BioMart and Bioconductor: a powerful link between biological databases and microarray data analysis. Bioinformatics 21(16): 3439–3440.

Edmond C, Shigaki T, Ewert S, Nelson MD, Connorton JM, Chalova V, Noordally Z, Pittman JK. 2009. Comparative analysis of CAX2-like cation transporters indicates functional and regulatory diversity. Biochemical Journal 418(1): 145–154.

Fernando DR, Lynch JP. 2015. Manganese phytotoxicity: new light on an old problem. Ann Bot 116(3): 313–319.

Ferrier S, Manion G, Elith J, Richardson K. 2007. Using generalized dissimilarity modelling to analyse and predict patterns of beta diversity in regional biodiversity assessment. Diversity and distributions 13(3): 252–264.

Fisher RA. 1930. The genetical theory of natural selection. Oxford,: The Clarendon press.

Fitzpatrick M, Mokany K, Manion G, Nieto-Lugilde D, Ferrier S. 2022. gdm: Generalized Dissimilarity Modeling. R package version 1.5.

Fitzpatrick MC, Keller SR. 2015. Ecological genomics meets community-level modelling of biodiversity: mapping the genomic landscape of current and future environmental adaptation. Ecology Letters 18(1): 1–16.

Folkertsma R, Charbonnel N, Henttonen H, Heroldova M, Huitu O, Kotlik P, Manzo E, Paijmans JLA, Plantard O, Sandor AD, et al. 2024. Genomic signatures of climate adaptation in bank voles. Ecology and Evolution 14(3): e10886.

Foy CD. 1984. Adaptation of plants to mineral stress in problem soils. Ciba Found Symp 102: 20–39.

Frachon L, Libourel C, Villoutreix R, Carrere S, Glorieux C, Huard-Chauveau C, Navascues M, Gay L, Vitalis R, Baron E, et al. 2017. Intermediate degrees of synergistic pleiotropy drive adaptive evolution in ecological time. Nat Ecol Evol 1(10): 1551–1561.

Garrison E, Kronenberg ZN, Dawson ET, Pedersen BS, Prins P. 2022. A spectrum of free software tools for processing the VCF variant call format: vcflib, bio-vcf, cyvcf2, hts-nim and slivar. PLOS Computational Biology 18(5): e1009123.

Garrison E, Marth G. 2012. Haplotype-based variant detection from short-read sequencing. arXiv preprint arXiv:1207.3907.

Goudet J, Jombart T. 2022. hierfstat: Estimation and tests of hierarchical F-statistics. R package version 0.5.

Gould B, McCouch S, Geber M. 2014. Variation in soil aluminium tolerance genes is associated with local adaptation to soils at the Park Grass Experiment. Molecular Ecology 23(24): 6058–6072.

Grossmann S, Bauer S, Robinson PN, Vingron M. 2007. Improved detection of overrepresentation of Gene-Ontology annotations with parent–child analysis. Bioinformatics 23(22): 3024–3031.

Guggisberg A, Liu X, Suter L, Mansion G, Fischer MC, Fior S, Roumet M, Kretzschmar R, Koch MA, Widmer A. 2018. The genomic basis of adaptation to calcareous and siliceous soils in Arabidopsis lyrata. Molecular Ecology 27(24): 5088–5103.

Hahn MW, Kern AD. 2005. Comparative genomics of centrality and essentiality in three eukaryotic protein-interaction networks. Molecular biology and evolution 22(4): 803–806.

Hamala T, Gorton AJ, Moeller DA, Tiffin P. 2020. Pleiotropy facilitates local adaptation to distant optima in common ragweed (Ambrosia artemisiifolia). PLoS Genet 16(3): e1008707.

Hendershot WH, Duquette M. 1986. A simple barium chloride method for determining cation exchange capacity and exchangeable cations. Soil science society of America journal 50(3): 605–608.

Huang Y, Tran I, Agrawal AF. 2016. Does genetic variation maintained by environmental heterogeneity facilitate adaptation to novel selection? The American Naturalist 188(1): 27–37.

Jumentier B, Caye K, Heude B, Lepeule J, François O. 2022. Sparse latent factor regression models for genome-wide and epigenome-wide association studies. Statistical Applications in Genetics and Molecular Biology 21(1): 20210035.

Karger DN, Conrad O, Böhner J, Kawohl T, Kreft H, Soria-Auza RW, Zimmermann NE, Linder HP, Kessler M. 2017. Climatologies at high resolution for the earth’s land surface areas. Scientific data 4(1): 1–20.

Kassambara A, Mund F. 2020. Factoextra: extract and visualize the results of multivariate data analyses. R package version 1.0.7.

Kinraide TB, Pedler JF, Parker DR. 2004. Relative effectiveness of calcium and magnesium in the alleviation of rhizotoxicity in wheat induced by copper, zinc, aluminum, sodium, and low pH. Plant and Soil 259(1-2): 201–208.

Kolde R. 2019. pheatmap: Pretty Heatmaps. R package version 1.0.12.

Lasky JR, Upadhyaya HD, Ramu P, Deshpande S, Hash CT, Bonnette J, Juenger TE, Hyma K, Acharya C, Mitchell SE. 2015. Genome-environment associations in sorghum landraces predict adaptive traits. Science advances 1(6): e1400218.

Lehmann B, Mackintosh M, McVean G, Holmes C. 2023. Optimal strategies for learning multi-ancestry polygenic scores vary across traits. Nature Communications 14(1).

Levene H. 1953. Genetic equilibrium when more than one ecological niche is available. The American Naturalist 87(836): 331–333.

Long S, Xie W, Zhao W, Liu D, Wang P, Zhao L. 2024. Effects of acid and aluminum stress on seed germination and physiological characteristics of seedling growth in Sophora davidii. Plant Signaling & Behavior 19(1): 2328891.

Lüdecke D. 2018. ggeffects: Tidy data frames of marginal effects from regression models. Journal of Open Source Software 3(26): 772.

Luqman H. 2021. Unravelling the neutral and adaptive genetic legacies of the past. ETH Zurich.

Luqman H, Wegmann D, Fior S, Widmer A. 2023. Climate-induced range shifts drive adaptive response via spatio-temporal sieving of alleles. Nature Communications 14(1): 1080.

Luu K, Bazin E, Blum MG. 2017. pcadapt: an R package to perform genome scans for selection based on principal component analysis. Molecular ecology resources 17(1): 67–77.

Mähler N, Wang J, Terebieniec BK, Ingvarsson PK, Street NR, Hvidsten TR. 2017. Gene co-expression network connectivity is an important determinant of selective constraint. PLoS genetics 13(4): e1006402.

McGee LW, Sackman AM, Morrison AJ, Pierce J, Anisman J, Rokyta DR. 2016. Synergistic Pleiotropy Overrides the Costs of Complexity in Viral Adaptation. Genetics 202(1): 285–295.

Michalet R, Gandoy C, Joud D, Pagès J-P, Choler P. 2002. Plant community composition and biomass on calcareous and siliceous substrates in the northern French Alps: comparative effects of soil chemistry and water status. Arctic, Antarctic, and Alpine Research 34(1): 102–113.

Moreno-Letelier A, Barraclough TG. 2015. Mosaic genetic differentiation along environmental and geographic gradients indicate divergent selection in a white pine species complex. Evolutionary ecology 29: 733–748.

Mostofa MG, Rahman MM, Ghosh TK, Kabir AH, Abdelrahman M, Khan MAR, Mochida K, Tran L-SP. 2022. Potassium in plant physiological adaptation to abiotic stresses. Plant Physiology and Biochemistry 186: 279–289.

Naciri R, Lahrir M, Benadis C, Chtouki M, Oukarroum A. 2021. Interactive effect of potassium and cadmium on growth, root morphology and chlorophyll a fluorescence in tomato plant. Scientific reports 11(1): 5384.

Nei M. 1987. Molecular evolutionary genetics. Columbia Univ.

Nocchi G, Whiting JR, Yeaman S. 2024. Repeated global adaptation across plant species. Proceedings of the National Academy of Sciences 121(52): e2406832121.

Obayashi T, Hibara H, Kagaya Y, Aoki Y, Kinoshita K. 2022. ATTED-II v11: a plant gene coexpression database using a sample balancing technique by subagging of principal components. Plant and Cell Physiology 63(6): 869–881.

Oksanen J. 2022. Vegan: community ecology package. R package version 2.6.4.

Orr HA. 1998. The population genetics of adaptation: The distribution of factors fixed during adaptive evolution. Evolution 52(4): 935–949.

Orr HA. 2000. Adaptation and the cost of complexity. Evolution 54(1): 13–20.

Paalsson A, Fior S, Widmer A. 2024. Trait evolution linked to climatic shifts contributes to adaptive divergence in an alpine carnation (Dianthus sylvestris). bioRxiv: 2024.2003. 2022.586106.

Panda SK, Khan MH. 2009. Growth, Oxidative Damage and Antioxidant Responses in Greengram (Vigna radiata L.) under Short-term Salinity Stress and its Recovery. Journal of Agronomy and Crop Science 195(6): 442–454.

Papatheodorou I, Fonseca NA, Keays M, Tang YA, Barrera E, Bazant W, Burke M, Füllgrabe A, Fuentes AM-P, George N. 2018. Expression Atlas: gene and protein expression across multiple studies and organisms. Nucleic acids research 46(D1): D246–D251.

Parisod C. 2022. Plant speciation in the face of recurrent climate changes in the Alps. Alpine Botany 132(1): 21–28.

Peiter E, Montanini B, Gobert A, Pedas P, Husted S, Maathuis FJ, Blaudez D, Chalot M, Sanders D. 2007. A secretory pathway-localized cation diffusion facilitator confers plant manganese tolerance. Proceedings of the National Academy of Sciences 104(20): 8532–8537.

Poulenard J, Podwojewski P 2006. Alpine soils. Encyclopedia of Soil Science.

Rajakaruna N. 2018. Lessons on evolution from the study of edaphic specialization. The Botanical Review 84: 39–78.

Rellstab C, Zoller S, Sailer C, Tedder A, Gugerli F, Shimizu KK, Holderegger R, Widmer A, Fischer MC. 2020. Genomic signatures of convergent adaptation to Alpine environments in three Brassicaceae species. Molecular Ecology 29(22): 4350–4365.

Rennison DJ, Peichel CL. 2022. Pleiotropy facilitates parallel adaptation in sticklebacks. Molecular Ecology 31(5): 1476–1486.

Russell L. 2024. Emmeans: estimated marginal means, aka least-squares means. R package version 1.10.3

Ryan P, Tyerman S, Sasaki T, Furuichi T, Yamamoto Y, Zhang W, Delhaize E. 2011. The identification of aluminium-resistance genes provides opportunities for enhancing crop production on acid soils. Journal of Experimental Botany 62(1): 9–20.

Sakaguchi S, Horie K, Ishikawa N, Nagano AJ, Yasugi M, Kudoh H, Ito M. 2017. Simultaneous evaluation of the effects of geographic, environmental and temporal isolation in ecotypic populations of Solidago virgaurea. New Phytologist 216(4): 1268–1280.

Shen Z, Wang J, Guan H. 1993. Effect of aluminium and calcium on growth of wheat seedlings and germination of seeds. Journal of plant nutrition 16(11): 2135–2148.

Sigel A, Sigel H. 1998. Metal ions in biological systems: CRC press.

Spichtig M, Kawecki TJ. 2004. The maintenance (or not) of polygenic variation by soft selection in heterogeneous environments. The American Naturalist 164(1): 70–84.

Storey JD, Bass AJ, Dabney A, Robinson D. 2022. qvalue: Q-value estimation for false discovery rate control. R package version 2.3.

Szklarczyk D, Franceschini A, Wyder S, Forslund K, Heller D, Huerta-Cepas J, Simonovic M, Roth A, Santos A, Tsafou KP. 2015. STRING v10: protein–protein interaction networks, integrated over the tree of life. Nucleic acids research 43(D1): D447–D452.

Takahashi K, Murayama Y. 2014. Effects of topographic and edaphic conditions on alpine plant species distribution along a slope gradient on Mount Norikura, central Japan. Ecological research 29: 823–833.

Tóth EG, Cseke K, Benke A, Lados BB, Tomov VT, Zhelev P, Kámpel JD, Borovics A, Köbölkuti ZA. 2023. Key triggers of adaptive genetic variability of sessile oak [Q. petraea (Matt.) Liebl.] from the Balkan refugia: outlier detection and association of SNP loci from ddRAD-seq data. Heredity 131(2): 130–144.

Troth A, Puzey JR, Kim RS, Willis JH, Kelly JK. 2018. Selective trade-offs maintain alleles underpinning complex trait variation in plants. Science 361(6401): 475–478.

Van Deurs S, Reutimann O, Luqman H, Lifshitz D, Mayzlish-Gati E, Alexander J, Fior S. 2025. Genomic Signatures of Adaptation Across a Precipitation Gradient From Niche Centre to Niche Edge. Molecular Ecology 34(6): e17696.

Vernham G, Bailey JJ, Chase JM, Hjort J, Field R, Schrodt F. 2023. Understanding trait diversity: the role of geodiversity. Trends in Ecology & Evolution 38(8): 736–748.

Wadgymar SM, DeMarche ML, Josephs EB, Sheth SN, Anderson JT. 2022. Local adaptation: causal agents of selection and adaptive trait divergence. Annual Review of Ecology, Evolution, and Systematics 53(1): 87–111.

Wagner GP, Kenney-Hunt JP, Pavlicev M, Peck JR, Waxman D, Cheverud JM. 2008. Pleiotropic scaling of gene effects and the ‘cost of complexity’. Nature 452(7186): 470–472.

Wagner GP, Zhang J. 2011. The pleiotropic structure of the genotype-phenotype map: the evolvability of complex organisms. Nature Reviews Genetics 12(3): 204–213.

Wang IJ, Bradburd GS. 2014. Isolation by environment. Molecular Ecology 23(23): 5649–5662.

Wang Y, Wu W-H. 2013. Potassium transport and signaling in higher plants. Annual review of plant biology 64(1): 451–476.

Wang Z, Liao BY, Zhang J. 2010. Genomic patterns of pleiotropy and the evolution of complexity. Proc Natl Acad Sci U S A 107(42): 18034–18039.

Wei T, Simko V. 2024. R package “corrplot”: Visualization of a Correlation Matrix. R package version 0.94.

Whiting JR, Booker TR, Rougeux C, Lind BM, Singh P, Lu M, Huang K, Whitlock MC, Aitken SN, Andrew RL, et al. 2024. The genetic architecture of repeated local adaptation to climate in distantly related plants. Nat Ecol Evol.

Williams AM, Ngo TM, Figueroa VE, Tate AT. 2023. The Effect of Developmental Pleiotropy on the Evolution of Insect Immune Genes. Genome Biology and Evolution 15(3).

Wollenberg Valero KC. 2020. Aligning functional network constraint to evolutionary outcomes. BMC Evolutionary Biology 20: 1–14.

Wood SN. 2011. Fast stable restricted maximum likelihood and marginal likelihood estimation of semiparametric generalized linear models. Journal of the Royal Statistical Society Series B: Statistical Methodology 73(1): 3–36.

Wos G, Arc E, Hülber K, Konečná V, Knotek A, Požárová D, Bertel C, Kaplenig D, Mandáková T, Neuner G. 2022. Parallel local adaptation to an alpine environment in Arabidopsis arenosa. Journal of Ecology 110(10): 2448–2461.

Yan J, Ye X, Song Y, Ren T, Wang C, Li X, Cong R, Lu Z, Lu J. 2023. Sufficient potassium improves inorganic phosphate-limited photosynthesis in Brassica napus by enhancing metabolic phosphorus fractions and Rubisco activity. The Plant Journal 113(2): 416–429.

Yanai T, Goto N, Goto J, Nonaka N, Shibata M. 2005. Morphological characteristics of nerve cells in the human general sensory system. Okajimas Folia Anat Jpn 82(2): 39–41.

Yeaman S. 2015. Local Adaptation by Alleles of Small Effect. American Naturalist 186: S74–S89.

Yeaman S. 2022. Evolution of polygenic traits under global local adaptation. Genetics 220(1).

Yeaman S, Chen Y, Whitlock MC. 2010. No effect of environmental heterogeneity on the maintenance of genetic variation in wing shape in Drosophila melanogaster. Evolution 64(12): 3398–3408.

Yeaman S, Jarvis A. 2006. Regional heterogeneity and gene flow maintain variance in a quantitative trait within populations of lodgepole pine. Proceedings of the Royal Society B: Biological Sciences 273(1594): 1587–1593.

Zhuo M, Sakuraba Y, Yanagisawa S. 2024. Dof1. 7 and NIGT1 transcription factors mediate multilayered transcriptional regulation for different expression patterns of nitrate transporter2 genes under nitrogen deficiency stress. New Phytologist 242(5): 2132–2147.

