## Supplementary Figures and Tables for "Soil heterogeneity and pleiotropy contribute to polygenic soil adaptation during postglacial range expansion in an alpine plant"

### Supplementary Tables and Figures

**Table S1**: A) Influence of soil variables and population genetic structure on genetic variation inferred by pRDA. B) Influence of soil variables, population genetic structure, and climatic variables on genetic variation inferred by pRDA. The proportion of explainable variance represents the total variance from the particular model (either with soil variables, population genetic structure or climatic variables as explanatory variable) divided by the variance explained by the full model. Note: G represents population allele frequencies as response variable.

A

| **Partial RDA models** | **Variance** | **$R^2$** | **Adjusted $R^2$** | **p value** | **Proportion of explainable variance** | **Proportion of total variance** |
|---|---|---|---|---|---|---|
| Full model: G ~ soil variables + population genetic structure | 26896 | 0.358 | 0.157 | 0.001 | 1 | 0.3577 |
| Soil model: G ~ soil variables \| (population genetic structure) | 14902 | 0.198 | 0.018 | 0.04 | 0.554 | 0.1982 |
| Population genetic structure model: G ~ population genetic structure \| (soil variables) | 7816 | 0.104 | 0.107 | 0.001 | 0.291 | 0.1039 |
| Total unexplained | 48301 | | | | | 0.6423 |
| Total variance | 75197 | | | | | 1 |

B

| **Partial RDA models** | **Variance** | **$R^2$** | **Adjusted $R^2$** | **p value** | **Proportion of explainable variance** | **Proportion of total variance** |
|---|---|---|---|---|---|---|
| Full model: G ~ soil variables + climatic variables + population genetic structure | 40994 | 0.55 | 0.2 | 0.001 | 1 | 0.5452 |
| Soil model: G ~ soil variables \| (population genetic structure + climatic variables) | 13616 | 0.18 | 0.01 | 0.146 | 0.33 | 0.1811 |
| Population genetic structure model: G ~ population structure/geography \| (soil variables + climatic variables) | 4979 | 0.07 | 0.08 | 0.001 | 0.12 | 0.0662 |
| Climatic model: G ~ climatic variables \| (soil variables + population genetic structure) | 14098 | 0.18 | 0.05 | 0.003 | 0.34 | 0.1875 |
| Total unexplained | 34203 | | | | | 0.4548 |
| Total variance | 75197 | | | | | 1 |

**Table S2**: Numbers of identified SNPs associated significantly with soil variables (Al, Ca, Co, Fe, K, Mg, Mn, Na, P, Sr, and pH) in LFMM2. Filtration steps are described in the Methods.

| | **LFMM2** | | | |
|---|---|---|---|---|
| **Variable** | **N of SNPs before filtrations** | **N of SNPs in genic regions** | **N of SNPs after allele frequency filtrations** | **N of SNPs after filtration for at least 2 SNPs/gene** |
| pH | 454 | 91 | 67 | 8 |
| Al | 128846 | 33114 | 7211 | 1380 |
| Ca | 3052 | 654 | 384 | 65 |
| Co | 2385 | 582 | 171 | 28 |
| Fe | 111731 | 28714 | 4908 | 937 |
| K | 210684 | 56737 | 6265 | 1093 |
| Mg | 60272 | 16172 | 3202 | 536 |
| Mn | 73511 | 18736 | 3037 | 556 |
| Na | 2281 | 607 | 153 | 25 |
| P | 7060 | 2062 | 251 | 32 |
| Sr | 21207 | 5549 | 1029 | 163 |

**Table S3**: Allelic turnover values for each soil variable in its associated candidate SNPs. Total numbers of associated SNPs are presented in Table 1.

| **Variable** | **Mean allelic turnover** | **Sd allelic turnover** |
|---|---|---|
| Al | 0.73 | 0.31 |
| Ca | 0.54 | 0.30 |
| Co | 0.42 | 0.18 |
| K | 0.97 | 0.51 |
| Mg | 0.67 | 0.36 |
| Mn | 0.53 | 0.22 |
| Na | 0.52 | 0.39 |
| P | 0.36 | 0.15 |
| pH | 0.30 | 0.13 |
| Sr | 0.53 | 0.21 |

**Table S4:** Overview of association of populations polygenic scores with the predicted probability of germination in high Al treatments. Seeds from these populations were germinated in two high Al treatments (presented here together). Predicted values for the probability of seed germination were extracted from a binomial generalized linear mixed effect regression model. Polygenic scores were calculated based on 240 Al-associated SNPs.

| Population | Polygenic score | Predicted probability of germination | CI 2.5% | CI 97.5% |
|---|---|---|---|---|
| 25 | 0.08 | 0.40 | 0.34 | 0.45 |
| 2 | 4.67 | 0.49 | 0.41 | 0.56 |
| 27 | 0.003 | 0.39 | 0.34 | 0.45 |
| 5 | 0.03 | 0.40 | 0.34 | 0.45 |
| 43 | 0.40 | 0.40 | 0.35 | 0.46 |
| 4 | 4.15 | 0.48 | 0.41 | 0.55 |

**Table S5:** Overview of number of gene-gene interactions inferred from STRING database for genes associated with metals (N = 76), genes associated with macronutrients (N = 53), and all candidate genes (N = 129). Mean number of interactions (N) with confidence intervals (CI) for randomly selected genes of *A. thaliana* was obtained from 1000 permutations.

| Gene categories | Mean N of interactions for candidate genes | Mean N of interactions for randomly selected genes | CI 2.5% for randomly selected genes | CI 97.5% for randomly selected genes |
|---|---|---|---|---|
| Genes associated with metals | 287.12 | 126.43 | 88.97 | 171.25 |
| Genes associated with macronutrients | 233.27 | 128.22 | 83.46 | 176.63 |
| All candidate genes | 265.24 | 127.33 | 99.17 | 157.57 |

**Table S6:** Overview of populations observed genetic heterozygosity ($H_o$) grouped into soil environments (identified in Figure 1d).

| Soil type | N of populations | Mean Ho |
|---|---|---|
| Soil environment 1 | 11 | 0.196 |
| Soil environment 2 | 5 | 0.193 |
| Soil environment 3 | 16 | 0.166 |
| Soil environment 4 | 7 | 0.181 |
| Soil environment 5 | 4 | 0.209 |

**Table S7:** Overview of *p*-values from Wilcoxon rank sum exact test for pairwise comparisons between soil types.

| | Soil environment 1 | Soil environment 2 | Soil environment 3 | Soil environment 4 |
|---|---|---|---|---|
| Soil environment 2 | 0.661 | | | |
| Soil environment 3 | 0.028 | 0.062 | | |
| Soil environment 4 | 0.193 | 0.185 | 0.193 | |
| Soil environment 5 | 0.457 | 0.185 | 0.017 | 0.040 |

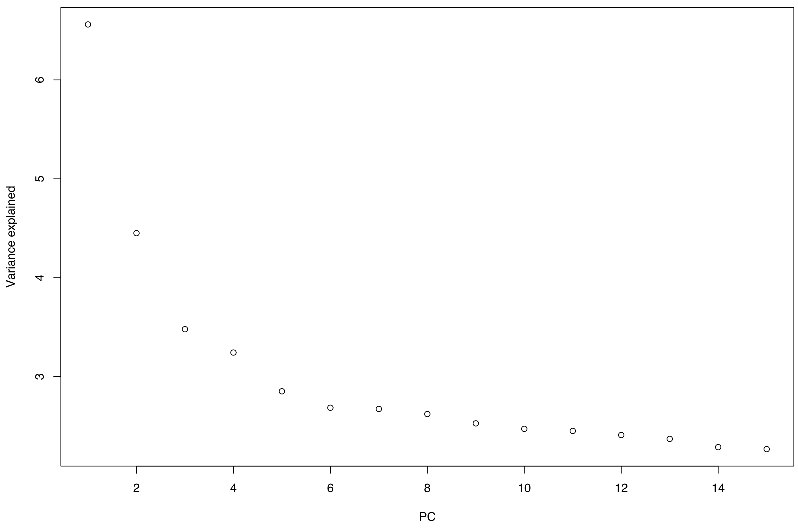


**Figure S1**: Determining K for environmental association analysis (LFMM2) with the scree plot. For details see Methods.

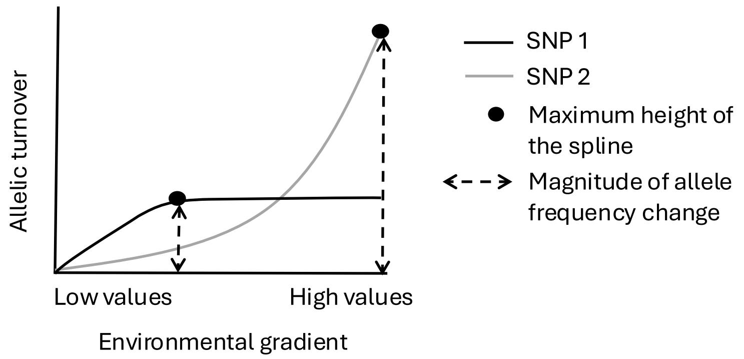

**Figure S2:** Illustration of the turnover functions to model allelic turnover along the environmental gradient. The maximum height of the GDM fitted spline indicates the total allelic turnover, i.e. magnitude of allele frequency change. The spline shape estimates how the rate of change in allele frequencies varies along the particular gradient. An example of two candidate SNPs: genetic variation in SNP 2 led to a higher allelic turnover along the gradient than genetic variation in SNP 1. The shape of the spline for SNP 1 indicates stronger response at low values than at higher values of the gradient. On the contrary, the spline for SNP 2 indicates strong response at high values of an environmental gradient. The figure was adapted from (Fitzpatrick & Keller, 2015; Storfer *et al.*, 2018).

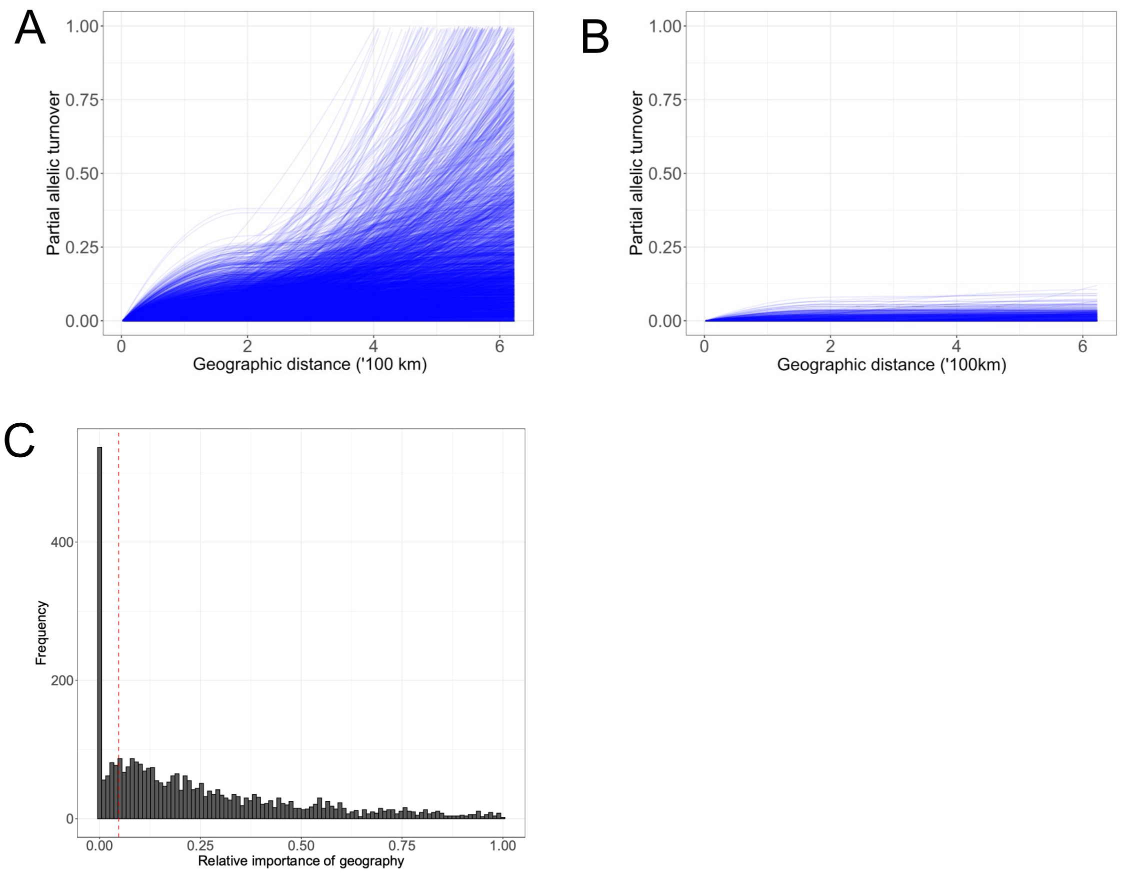


**Figure S3:** Allelic turnover relationship with geographic distance. A) Allelic turnover function for 3763 SNPs, that showed significant allelic turnover associated with any of the soil variables or geographic distance in GDM. Here, we specifically visualised partial allelic turnover in relation to geographic distance. B) Allelic turnover function for 814 candidate SNPs after removing SNPs that had a relatively strong association with geographic distance compared to soil variables. C) Relative importance of geography from GDM; the dashed line indicates 25% of the distribution (<= 0.047), we kept only SNPs below that threshold.

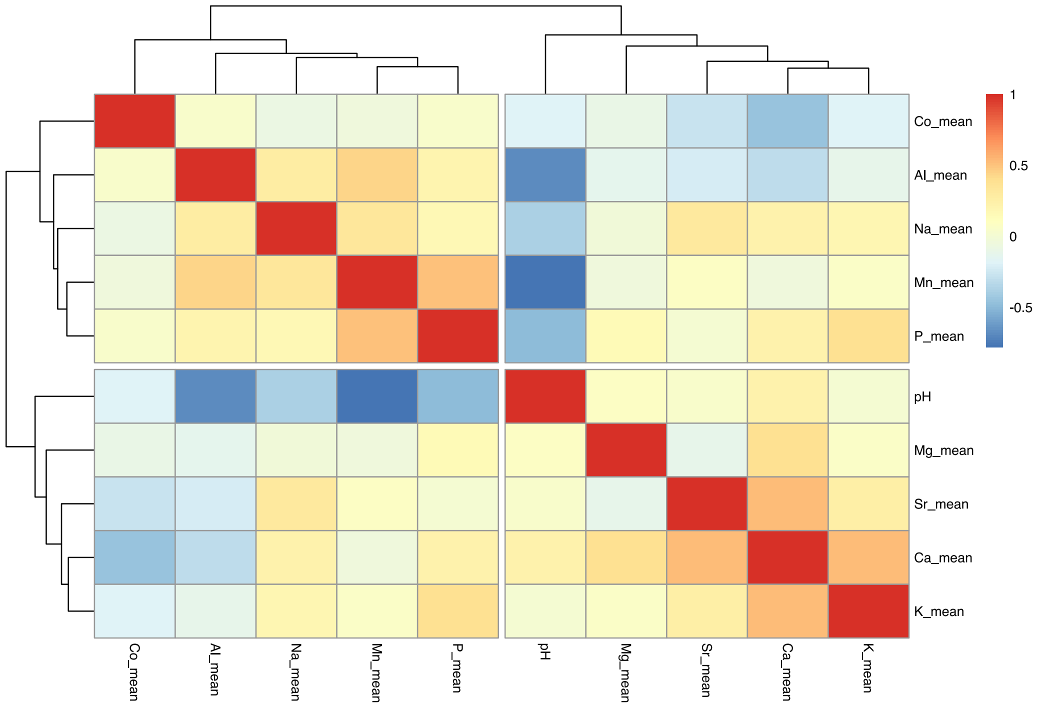


**Figure S4:** Correlation of exchangeable cations in the soil. Clustering based on Spearman's correlation coefficients among exchangeable cations indicates two groups of elements – "macronutrients" (Ca, K, Mg, pH, and Sr) and "metals" (Al, Co, Mn, Na, and P).

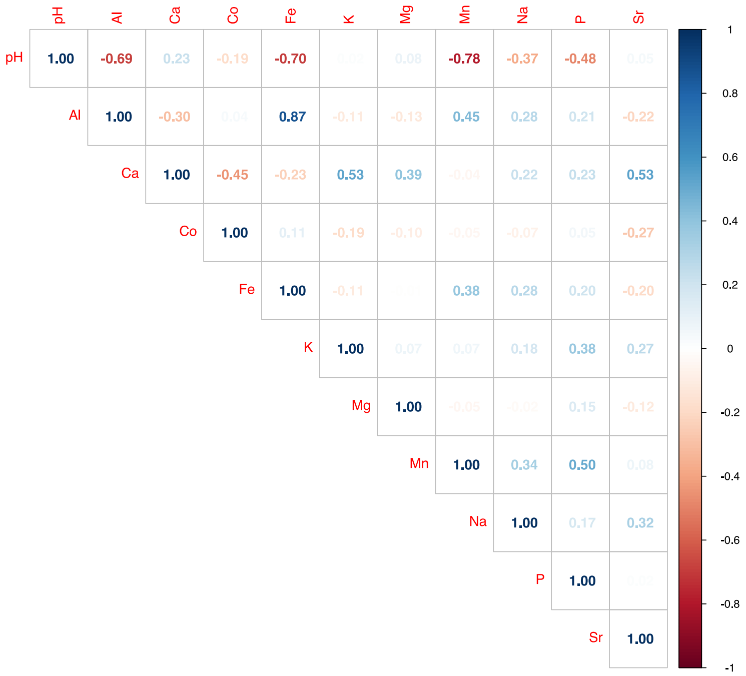


**Figure S5**: Pairwise Spearman's correlation coefficients among exchangeable concentrations and pH.

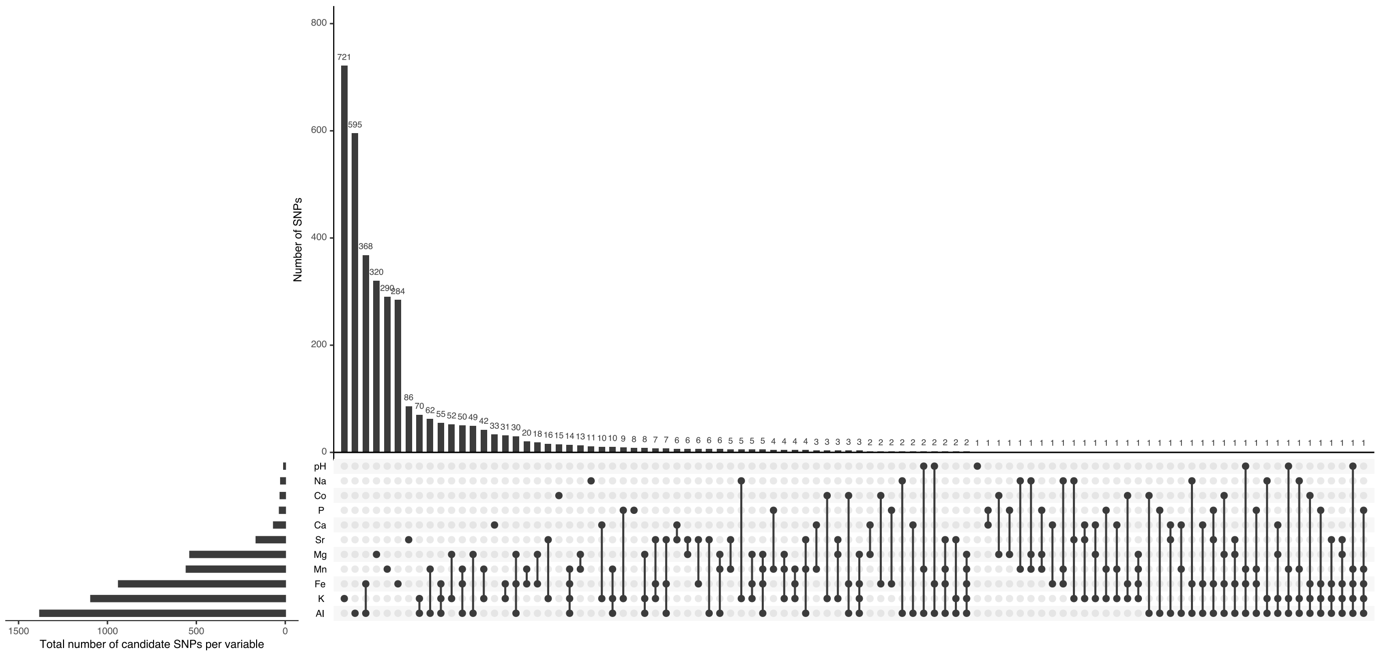


**Figure S6:** Intersection plot of significantly associated SNPs with pH and exchangeable concentrations identified by LFMM2.

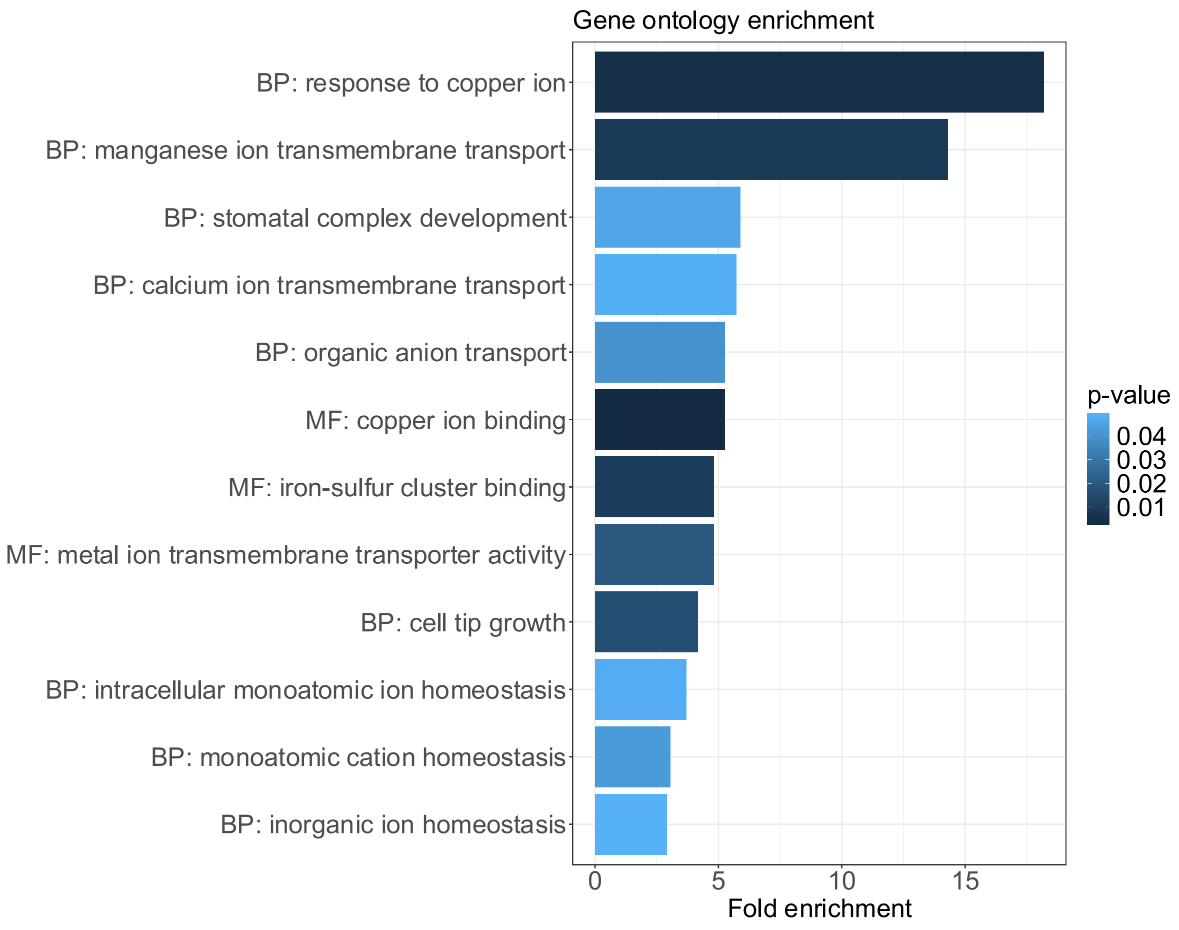

**Figure S7:** Gene ontology (GO) enrichment of biological processes (BP) and molecular functions (MF) of 129 candidate genes. Selection of relevant BP and MF for soil adaptation from the complete list of significantly enriched terms (Data S7). Fold enrichment indicates how overrepresented is a particular GO term in a given set of genes compared to a background of *A. thaliana* genes. All terms were significantly enriched ($p < 0.05$, Fisher's exact test).

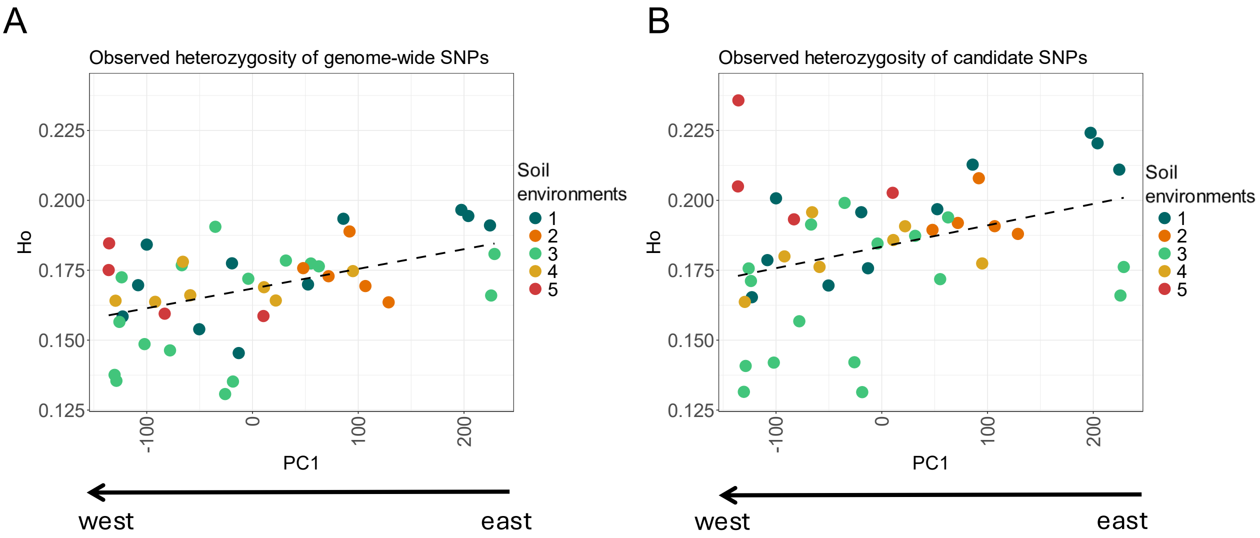


**Figure S8:** Genetic diversity of 43 populations along range expansion in the Alps. Observed heterozygosity ($H_o$) of genome-wide exonic SNPs (A) and candidate SNPs (B) in populations along range expansion. Note: x-axis with principal component 1 scores represents the expansion axis from eastern to western Alps. Colours denote the assignement of populations to the soil environments from Figure 1d.

References

**Fitzpatrick MC, Keller SR. 2015.** Ecological genomics meets community-level modelling of biodiversity: mapping the genomic landscape of current and future environmental adaptation. *Ecology Letters* **18**(1): 1-16.

**Storfer A, Patton A, Fraik AK. 2018.** Navigating the Interface Between Landscape Genetics and Landscape Genomics. *Front Genet* **9**: 68.
