## Supplementary material for "Soil heterogeneity and pleiotropy contribute to polygenic soil adaptation during postglacial range expansion in an alpine plant": Overview_supplementary_data.docx

Data S1: Population environmental data (exchangeable cations [mg/kg], pH, and climatic data) and geographical locations

Data S2 A-K: Significantly associated SNPs with at least one soil variable (pH, Al, Ca, Co, Fe, K, Mg, Mn, Na, P, and Sr) identified by LFMM2.

Data S3 A-K: Refined list of significantly associated SNPs with at least one soil variable (pH, Al, Ca, Co, K, Mg, Mn, Na, P, and Sr) identified by LFMM2.

Data S4: List of 814 candidate SNPs identified by GDM.

Data S5: Overview of experimental populations with exchangeable cations of original populations, individual seed germination rates, and mean population germination rates.

Data S6: List of 129 candidate genes with descriptions.

DataS7: Gene ontology enrichment of 129 candidate genes.

Data S8: Proxies for pleiotropy of candidate genes.
